# Heme-Mediated Selection of Encapsulated *Streptococcus pneumoniae* in the Lungs by Oxidative Stress

**DOI:** 10.1101/2023.11.14.567109

**Authors:** Babek Alibayov, Ana G. Jop Vidal, Landon Murin, Anna Scasny, Kenichi Takeshita, Komal Beeton, Kristin S. Edwards, Tracy Punshon, Brian P. Jackson, Larry S. McDaniel, Jorge E. Vidal

## Abstract

*Streptococcus pneumoniae* (the pneumococcus) causes cytotoxicity and encapsulates within the lung parenchyma, leading to pneumococcal pneumonia. However, the underlying mechanisms remain unclear and likely involve multiple bacterial and host factors.

We investigated the selection process of encapsulated pneumococci, a critical factor in lung damage during pneumococcal pneumonia.

Our study revealed that pneumococci initially lack capsules but re-encapsulate upon reaching the alveoli. This process is driven by *S. pneumoniae*-derived hydrogen peroxide (Spn-H₂O₂), which oxidizes lung hemoglobin, leading to heme release and polymerized hemoglobin formation. Physiologically relevant levels of heme were found to promote the selection of encapsulated bacteria. Furthermore, encapsulation protects bacteria from intracellular heme toxicity, a defense absent in non-encapsulated strains. Ultrastructural analysis demonstrated interactions between hemoglobin and both encapsulated and non-encapsulated pneumococci in human sputum.

These findings reveal a critical connection between oxidative stress-mediated lung damage and the selection of encapsulated pneumococci, suggesting potential therapeutic avenues by targeting these oxidative processes.

## Introduction

*Streptococcus pneumoniae* (Spn) strains colonize the lungs causing pneumonia, a global disease that kills more than a million individuals every year.^1^ Invasion of bronchi and terminal bronchiole allows pneumococci to reach respiratory bronchiole, alveolar ducts, and alveoli, where pneumococci have access to the alveolar-capillary network, a large surface area containing ∼300 billion of capillaries and where the exchange of CO by O occurs.^2, 3^

In the lungs where molecular oxygen (O_2_) is available, Spn uses a fermentative metabolism to produce ATP by oxidizing glucose to acetate.^4, 5, 6^ Spn strains encode an incomplete set of genes for enzymes of the Krebs cycle, or proteins of the respiratory chain. ATP production includes the oxidation of pyruvate by the enzyme pyruvate oxidase (SpxB), releasing H_2_O_2_ as a byproduct.^4, 5, 6^ In the absence of O_2_, pyruvate is converted to L-lactate, whereas, in the presence of oxygen, L-lactate is oxidized back to pyruvate, via lactate oxidase (LctO), producing H_2_O_2_ as a byproduct.^4, 5^

Besides components of the glycolytic pathway and atmospheric O_2_, the alveolar-capillary network is a source of erythrocytes and hemoglobin. We recently demonstrated that hemoglobin enhances *in vitro* growth of Spn strains and upregulates expression of *spxB*.^7, 8^ H_2_O_2_ produced by the pneumococcus oxidizes oxy-hemoglobin (Fe^2+^) from different mammals, including humans, to produce the non-oxygen-binding form of hemoglobin known as met-hemoglobin (Hb-Fe^3+^).^9, 10^ The oxidation of hemoglobin by Spn produced toxic species, including labile heme^11^ and a hemoglobin-derived tyrosyl radical [Fig. 1A, (^•^Hb-Fe^4+^)].^12^ These toxic products are implicated in the pathophysiology of human diseases characterized by intravascular hemolysis, such as sickle cell disease (SCD), and in inflammatory conditions, such as sepsis.^13, 14, 15^

**Fig. 1.**
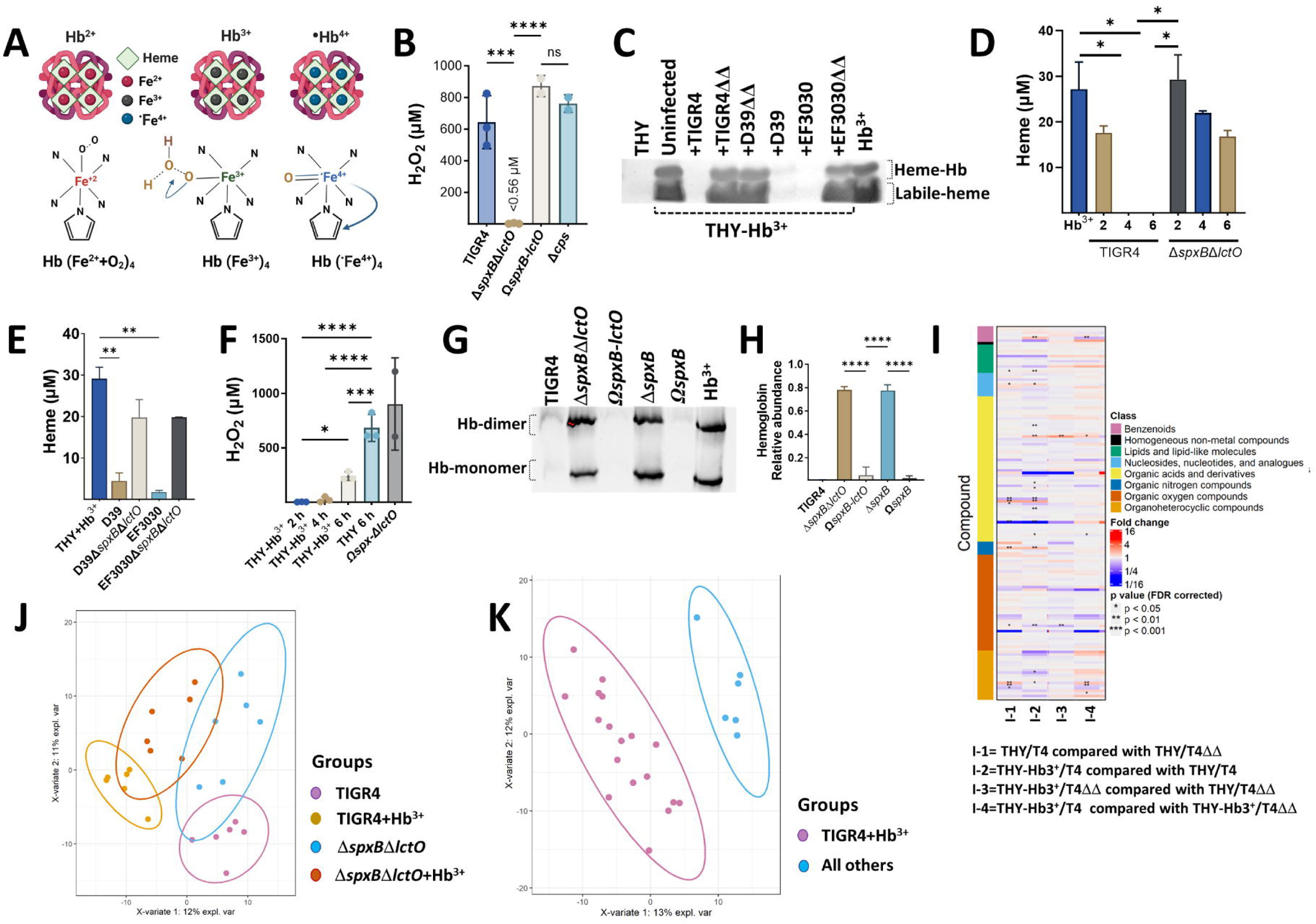
Oxidation of hemoglobin by Spn creates metabolites that contributes to oxidative stress. (A) Schematic representation of oxy-hemoglobin (Hb^2+^), met-hemoglobin (Hb^3+^), and ferryl-hemoglobin (•Hb^4+^). (B) Spn was cultured in THY medium at 37°C for 6 h. Supernatants were collected, and H₂O₂ concentration was quantified using Amplex Red (LOD < 0.56 µM). (C) THY added with met-hemoglobin (THY-Hb^3+^) was inoculated with Spn or left uninfected. Supernatants, THY alone, or Hb (10 µM) were analyzed by native gel electrophoresis with in-gel heme staining. (D & E) Heme, or (F) H₂O₂, was quantified in supernatants from (C) using the QuantiChrom Heme Assay or Amplex Red assays, respectively. (G) Supernatants from cultures in (C) and Hb^3+^ (10 µM) were analyzed by Western blot with a mouse monoclonal anti-β-hemoglobin antibody. (H) Densitometric analysis of Hb monomer relative to control Hb^3+^. Data in panels B, D-F, and H represent mean ± SEM from ≥ 3 independent experiments. Statistical significance was determined by one-way ANOVA with Dunnett’s post hoc test (*p < 0.05, **p < 0.001, ***p < 0.0005, ****p < 0.0001). (I) Heatmap depicting pairwise comparisons between groups for identified molecules. Each row represents a chemical compound, and each column represents a comparison. Asterisks indicate statistical significance by Mann-Whitney U test. The color bar indicates the chemical superclass. (J) PCA biplot showing the first two principal components with group membership and normal data ellipses. (K) PLS-DA biplots of the first two components, showing group membership and normal data ellipses.

This study demonstrates that hydrogen peroxide produced by Spn (Spn-H₂O₂) oxidizes hemoglobin within the alveolar-capillary network, driving the differentiation of encapsulated pneumococci within the alveolar space. This oxidation process leads to hemoglobin polymerization, which disrupts the alveolar epithelium and is closely associated with pneumococci. Ultrastructural analysis of human sputum revealed interactions between hemoglobin and both encapsulated and non-encapsulated pneumococci. Encapsulated, H₂O₂-producing pneumococci exhibit resistance to physiological levels of free heme, unlike their unencapsulated, H₂O₂-deficient counterparts, which are more susceptible to heme-mediated toxicity. These findings highlight the capsule’s role as a protective barrier against heme-mediated toxicity and suggest its potential as a therapeutic target for pneumococcal pneumonia.

## Results

### Spn oxidizes hemoglobin generating molecules of oxidative stress

Reactive oxygen species (ROS) can contribute to heme degradation by damaging the protoporphyrin IX ring (Fig. 1A)^16, 17, 18^. To investigate Spn-H_2_O_2_ involvement in oxidative stress and hemoglobin oxidation, we assessed H_2_O_2_ in Spn cultures. Wild-type Spn strains (TIGR4, D39, and EF3030) generated substantial H_2_O_2_ (>650 µM) at 6 h post-inoculation in THY medium (Fig. 1B, and Fig. S1A) whereas the isogenic H_2_O_2_-defective Δ*spxB*Δ*lctO* mutants did not (Fig. 1B and Fig. S1A). To further explore this, Spn and Δ*spxB*Δ*lctO*, strains were cultured in THY supplemented with met-hemoglobin (THY-Hb^3+^). Heme levels in supernatants were quantified using an in-gel heme staining (Fig. S1B), and a Quantichrom assay.^19, 20^ Compared to the uninfected control, heme levels in Spn cultures were significantly reduced or undetectable within 6 hours (Fig. 1C-E, Fig. S1C). In contrast, THY-Hb^3+^ cultures of Δ*spxB*Δ*lctO* strains yielded >16.82 µM of total heme at 6 h (Fig. 1C-E). This heme degradation correlated with Spn-mediated pseudoperoxidase activity, as Spn-H_2_O_2_ levels were undetectable in THY+Hb^3+^ cultures 2 or 4 h post-inoculation (Fig. 1F).

To confirm hemoglobin oxidation, we cultured TIGR4 and TIGR4Δ*spxB*Δ*lctO* in THY or in THY-Hb^3+^ for two h. Subsequent metabolomic analysis using partial least squares discriminant analysis (PLS-DA) revealed distinct metabolic profiles between groups (Fig. 1J). The largest biological variation was observed between TIGR4 cultures grown in THY-Hb^3+^ (component 2, 10% variance; AUC=0.9542) (Fig. 1K). A heatmap summarizing metabolites with at least a 2-fold change and FDR p-value <0.05 is presented (Fig. 1J). Compared to THY cultures, TIGR4 grown with THY-Hb^3+^ exhibited increased levels of metabolites indicative of hemoglobin oxidation, such as creatinine, isoleucine, leucine, guanosine, putrescine, and tyrosyl. Conversely, aspartic acid, guanine, and oxoproline levels decreased (Table S1). Interestingly, TIGR4 cultures with H_2_O_2_(THY-Hb^3+^) showed increased levels of oxidative stress markers,^21^ like methionine sulfoxide (MetO), oleamide, and putrescine (Table S1) compared to Δ*spxB*Δ*lctO* mutant cultures. Additionally, methionine (precursor to MetO) and pyruvic acid (SpxB substrate) were downregulated (Table S1). These findings collectively suggest that Spn-derived H_2_O_2_ oxidizes hemoglobin and generating metabolites associated with oxidative stress.

### Spn oxidation and polymerization of hemoglobin favor the selection of encapsulated bacteria

Oxidation crosslinks hemoglobin globins, potentially leading to precipitation.^22, 23^ Consistent with this, Spn cultures in hemoglobin-supplemented media (THY-Hb^3+^) exhibited reduced levels of soluble α and β globins upon oxidation (Fig. 1G-H, Fig. S1D and, S1E). In contrast, H_2_O_2_-deficient mutants displayed normal hemoglobin monomer and dimer levels in THY-Hb^3+^, similar to uninfected controls (Fig. 1G-H, Fig. S1D).

We next investigated the structural localization of the polymerized hemoglobin aggregates. Confocal microscopy analysis of biofilms from Spn cultures grown in THY-Hb^3+^ included capsule, hemoglobin, and DNA staining. Z-stack projections and their 3D reconstruction revealed bacteria associated with hemoglobin (hereafter Spn-Hb^+^) as early as 2 h post-inoculation, with increasing abundance over time (Fig. 2A, Fig. S2A-B). Structures suggestive of hemoglobin polymers were also observed (Fig. S2C and S2D). Additionally, Spn-Hb^+^ bacteria were positive for staining with anti-alpha chain hemoglobin antibodies (Not shown).

**Fig. 2.**
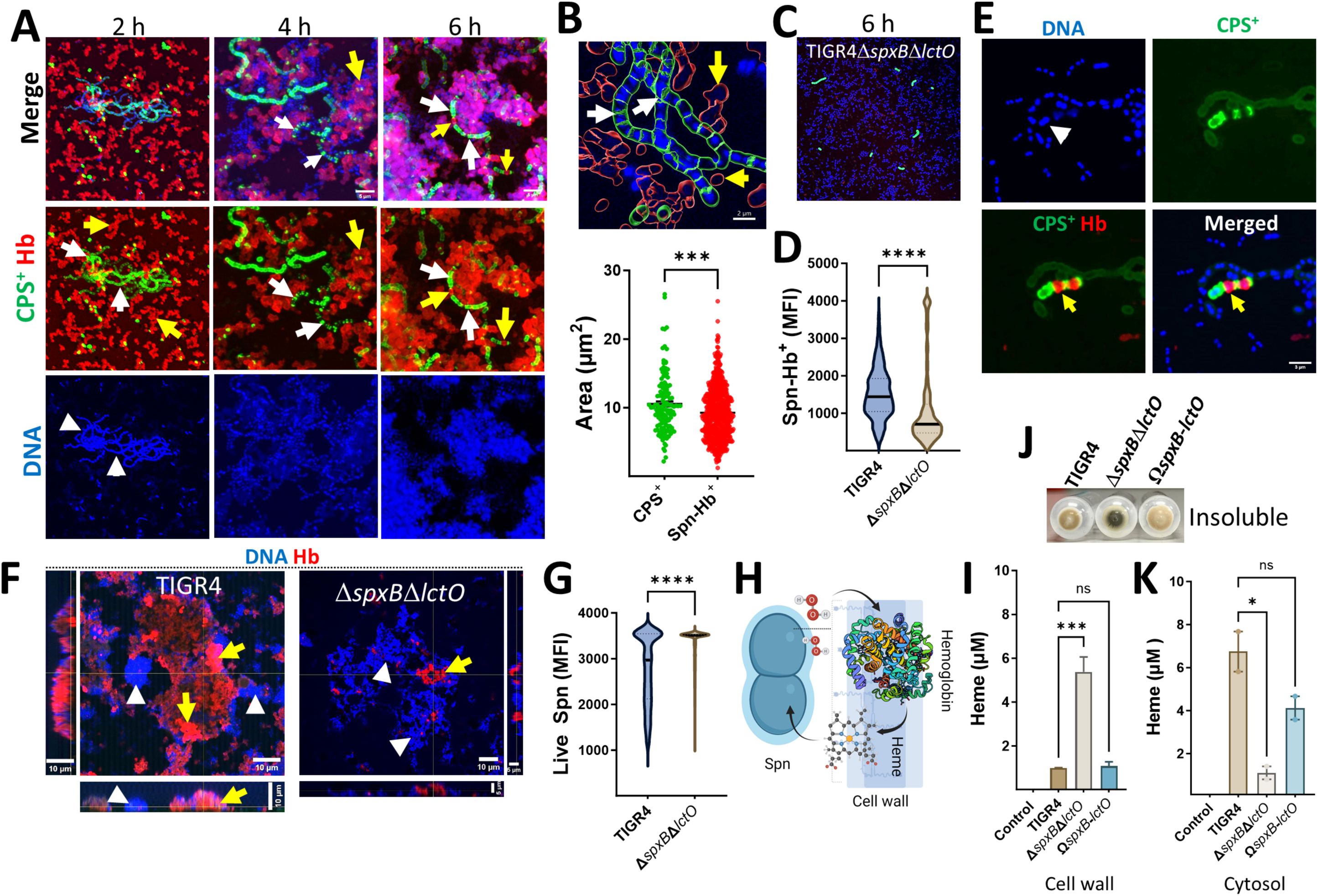
Oxidation of hemoglobin by Spn selects for hemoglobin-binding (Spn-Hb^+^) and encapsulated pneumococci. TIGR4 (A, B, E) or TIGR4Δ*spxB*Δ*lctO* (C) were incubated with Hb^3+^ and analyzed. Biofilms (A, C) or planktonic bacteria (E) were stained for capsule (green), Hb (red), and DNA (blue). Images were acquired by confocal microscopy and represent z-stack projections. Arrows: white, CPS^+^ bacteria; yellow, Spn-Hb^+^ bacteria; arrowheads, DNA. (B) Area occupied by CPS^+^ and Spn-Hb^+^ bacteria was quantified using Imaris software. The top panel shows masks used for quantification. (D) Minimum fluorescent intensity (MFI) of Spn-Hb bacteria (red channel) in TIGR4 (A) or TIGR4Δ*spxB*Δ*lctO* (C) cultures incubated for 6 h was measured with Imaris software. (F) TIGR4 or TIGR4Δ*spxB*Δ*lctO* were incubated with Hb for 6 h. Planktonic bacteria were collected, stained for Hb and DNA, and analyzed by confocal microscopy. XY, XZ (bottom), and YZ (side) optical sections are shown. (G) MFI of the DAPI channel (live Spn) was measured using Imaris software. (H) Schematic diagram depicting the hypothesized mechanism: Hb binds a cell wall receptor, is oxidized by Spn-H_2_O_2_, releasing heme that enters the cytosol. (J, K) Spn was incubated with Hb for 4 h. Cellular fractionation was performed, and the insoluble fraction is shown in (J). Heme was quantified from the (I) cell wall fraction or the (K) cytosolic fraction using the QuantiChrom Heme Assay. Data (B, D, G, I, K) are presented as mean ± SEM of at least three independent experiments. Statistical analysis: Student’s t-test (B, D, G) or one-way ANOVA with Dunnet’s test for multiple comparisons (I, K). ns, not significant; *, p < 0.05; ***, p < 0.005; **, p < 0.0001.

Confocal microscopy analysis revealed that a significant proportion of Spn-Hb^+^ were non-viable. Quantitative analysis of images showed only 12% of Hb^+^-associated bacteria colocalized with DNA (a viability marker), compared to 76% of capsular polysaccharide-positive (CPS^+^) pneumococci (Fig. 2B and not shown). Furthermore, the DNA area of Spn-Hb^+^ bacteria was significantly smaller than that of CPS^+^ pneumococci (Fig. 2B), suggesting DNA degradation due to oxidative stress. Notably, cultures with the H_2_O_2_-deficient Δ*spxB*Δ*lctO* strain showed a significant decrease in both dead Spn-Hb^+^ bacteria (Fig. 2C and 2D) and CPS^+^ pneumococci (Fig. 2C). Similar trends were observed in planktonic cultures, with wild-type exhibiting increased Spn-Hb^+^ and reduced viability, while the *ΔspxBΔlctO* mutant showed the opposite (Fig. 2E-G, S2F, S3A-C). These findings suggest that oxidative products of hemoglobin, including heme (assessed in a later section), contribute to bacterial toxicity.

The exclusive presence of Hb^+^ bacteria in wild-type Spn cultures suggests hemoglobin binds to the cell wall, followed by H_2_O_2_-mediated oxidation (Fig. 2H). To investigate heme location, we quantified heme levels in cytosolic, insoluble and cell wall fractions of Spn grown with THY-Hb^3+^. Heme was detected in the insoluble fraction (Fig. 2J) and significantly increased in the cell wall fraction of H_2_O_2_-deficient Δ*spxB*Δ*lctO* mutant compared to wild-type Spn (Fig. 2I). Conversely, the cytosolic fraction of H_2_O_2_-producing Spn contained significantly more heme compared to the Δ*spxB*Δ*lctO* mutant, indicating that hemoglobin oxidation facilitates heme uptake (Fig. 2K). These findings demonstrate that Spn-H_2_O_2_-mediated hemoglobin oxidation contributes to bacterial killing, hemoglobin polymerization, and the selection of encapsulated pneumococci, potentially enhancing survival during infection.

### Spn-H_2_O_2_ oxidation of hemoglobin contributes to mortality in a mouse model of pneumococcal disease

Since hemorrhage, a hallmark of lobar pneumococcal pneumonia (also known as red hepatization)^24^, involves hemoglobin, we employed a mouse model to assess the contribution of H_2_O_2_-mediated hemoglobin oxidation to pneumococcal pneumonia severity. The mouse model mimics the natural course of infection, where Spn is aspirated into the lower airways, facilitating colonization in the bronchi, bronchioles, and/or alveoli (Fig. 3A). Since the capsule *in vivo* protects Spn from the immune response^2, 25, 26, 27^, and mutations in *spxB* affect capsule production in some strains^28^, we quantified CPS^+^ bacteria in strains used to inoculate animals (Fig. 3B). The majority of TIGR4, TIGR4Ω*spxB*-*lctO,* D39, and D39Δ*spxB*Δ*lctO* (>84.1%) expressed the capsule, whereas the capsule-deficient TIGR4Δ*cps* mutant and TIGR4Δ*spxB*Δ*lctO* exhibited significantly lower capsular polysaccharide-positive (CPS⁺) populations (<28.8%), compared to TIGR4 (Fig. 3C). Similar results were obtained using an ELISA approach (Data not shown). Deletion of *spxB* and *lctO* affected production of capsule in TIGR4 but not in D39, indicating strain-specific effects

**Fig. 3.**
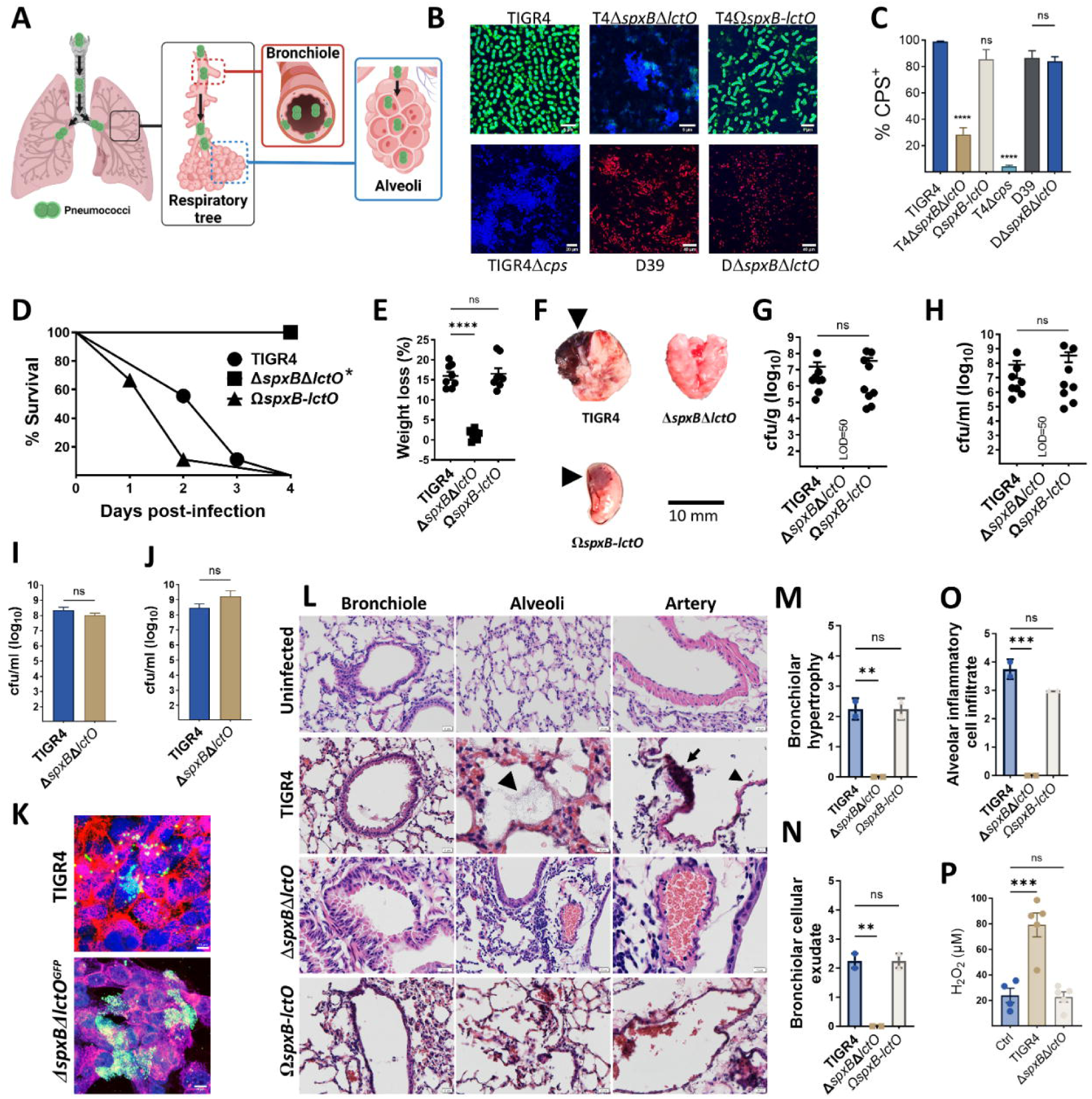
Pneumococci lacking capsule and hydrogen peroxide are rapidly cleared from the lungs. (A) Schematic representation of the murine pneumococcal pneumonia model. (B) Confocal micrographs of capsule-stained pneumococci. (C) Quantification of capsule production (percentage capsule-positive bacteria). (D-H) Groups of eight Balb/c mice were intranasally inoculated with approximately ∼10^7^ colony-forming units (CFU) of the indicated strain. Survival (D) and weight loss (E) were monitored. At endpoint, lung tissues were harvested, imaged (F), homogenized to quantify bacterial burden (G), and processed for histological analysis (L). Blood was collected for bacterial quantification (H). (I) Human alveolar A549 (J) or mouse alveolar MLE-12 (J) cells were infected with TIGR4 or TIGR4^GFP^Δ*spxB*Δ*lctO* for 6 hours, followed by bacterial enumeration. (K) Confocal microscopy of membrane-, DNA-, and capsule-stained A549 cells infected with pneumococci. (M-O) Lung sections were subjected to histopathological analysis, and scores for bronchiolar hypertrophy (M), bronchiolar cellular exudate (N), and alveolar inflammatory cell infiltrate (O) were assigned by a blinded pathologist. (P) Hydrogen peroxide levels in lung homogenates from mock-infected or pneumococcus-infected mice. Data are presented as mean ± SD. One-way ANOVA with Dunnett’s multiple comparisons test was used for (C, E, G, H, M-P), and Mann-Whitney test for (I, J). Survival analysis was performed using the log-rank Mantel-Cox test. *p<0.0001, **p<0.01, ***p<0.001, ****p<0.0001. ns: not significant.

Balb/c mice were then intranasally infected with wild-type strains (TIGR4 or D39) or H_2_O_2_-deficient mutants. Mice infected with TIGR4, complemented TIGR4Ω*spxB*-*lctO,* or D39 succumbed within 96 h post infection (Fig. 3D and Supplementary Fig. 4A, 5A). In contrast. mice infected with TIGR4Δ*spxB*Δ*lctO* survived at least 96 h, and a significant proportion of mice infected with D39Δ*spxB*Δ*lctO* also survived (Fig. 3D, and Supplementary Fig. 5A). Notably, D39-infected mice succumbed 3.5 times faster than the D39Δ*spxB*Δ*lctO* group (not shown). Weight loss, another indicator of disease severity, was also monitored. Animals infected with H_2_O_2_-producing strains displayed significantly greater body weight reduction at the time of euthanasia compared to their isogenic Δ*spxB*Δ*lctO* mutant counterpart (Fig. 3E, Supplementary Fig. 4B, and 5B).

Lung pathology in mice infected with TIGR4, or TIGR4Ω*spxB*-*lctO* resembled human lobar pneumonia, exhibiting hemorrhage with focal sub-lobar lesions^24^ (Fig. 3F). In contrast, lungs from mice infected with H_2_O_2_-deficient strain appeared normal (Fig. 3F). This observation aligned with bacterial burden data: the median lung density of TIGR4 reached 2.3x10^7^ cfu/g, while TIGR4Ω*spxB*-*lctO* reached 1.4x10^8^ cfu/g (Fig. 3G, and Supplementary Fig. 4C). Importantly, the H_2_O_2_-deficient strain was completely cleared from the lungs by 96 h post-infection (Fig. 3G). Consistent with lung colonization, pneumococci were detected in the blood of mice infected with TIGR4, or TIGR4Ω*spxB*-*lctO,* (3.4x10^6^ and 2.8 x10^6^ cfu/ml, respectively) (Fig. 3H) but not in those infected with H_2_O_2_ - deficient strain (Fig. 3H).

To investigate if the rapid clearance of the H_2_O_2_-deficient strain from the lung was due to decreased adhesion, we infected human (A549), and mouse (MLE-12), alveolar type I pneumocytes with TIGR4 or a GFP-tagged H_2_O_2_-deficient strain (TIGR4Δ*spxB*Δ*lctO^GFP^*). Bacterial burden, quantified by both culture and confocal microscopy, demonstrated comparable levels for both strains in both human and mouse pneumocytes (Fig. 3I, 3J and 3K).

### Histopathological examination of lung tissue revealed extensive damage induced by H_2_O_2_-producing pneumococci

Mice infected with H_2_O_2_-producing strains, TIGR4 and D39, exhibited severe lung pathology characterized by bronchiolar epithelial detachment, fibrotic alveolar changes, alveolar hemorrhage, and pneumococcal infiltration (Fig. 3L, Supplementary Fig. 5D). Alveolar artery endothelial damage was also evident in these mice. In contrast, H_2_O_2_-deficient strains, TIGR4*ΔspxBΔlctO* and D39*ΔspxBΔlctO*, induced milder lung pathology with primarily inflammatory cell infiltration and intimal/medial thickening of alveolar arteries, but without endothelial cytotoxicity (Fig. 3L, Supplementary Fig. 5D). While TIGR4*ΔspxBΔlctO* caused minimal bronchiolar hemorrhage and no fibrosis or cytotoxicity, D39*ΔspxBΔlctO* induced localized inflammation and some alveolar fibrosis without endothelial damage (Fig. 3L and Supplementary Fig. 4E, and 5D). Histopathological scoring revealed significantly higher scores for bronchiolar hypertrophy, cellular exudate, and alveolar inflammatory cell infiltrate in lungs from mice infected with TIGR4 or complemented TIGR4*ΩspxB-lctO* compared to those infected with TIGR4*ΔspxBΔlctO* (Fig. 3M-O). These findings indicate that H_2_O_2_-producing pneumococci induce substantial cytotoxic damage across multiple lung cell types.

### Increased reactive oxygen species in the lungs of mice infected with H_2_O_2_*-*producing pneumococci

To assess H_2_O_2_ production by pneumococci *in vivo*, we measured ROS levels in lung tissue. Mice infected with wild-type TIGR4 for 24 h exhibited significantly elevated lung ROS levels compared to uninfected mice or those infected with the H_2_O_2_-deficient Δ*spxB*Δ*lctO* strain (Fig. 3P), confirming Spn-H_2_O_2_ production *in vivo*. While host immune cells such as neutrophils likely contribute to the overall ROS levels, a human lung cell model was used to directly evaluate pneumococcus-derived ROS. In this model, intracellular ROS in alveolar A549 cells infected with TIGR4 was significantly elevated, mirroring the effect of menadione, a positive redox control (Supplementary Fig. 6A-B)^29^. Confocal microscopy confirmed intracellular pneumococci in both TIGR4 and *ΔspxBΔlctO* infected cells (Supplementary Fig. 6C), suggesting H_2_O_2_ from TIGR4 contributed to the ROS rise. As H_2_O_2_ exposure can damage cellular structures, we examined potential damage to tubulin and GAPDH, proteins known to be sensitive to H_2_O_2_ ^30, 31^. Western blotting revealed significant degradation of β-tubulin and GAPDH in human alveolar cells infected with TIGR4 but not in those infected with the H_2_O_2_-deficient *ΔspxBΔlctO*, while cytosolic catalase levels remained unchanged (Supplementary Fig. 6D-F). These findings demonstrate that Spn-H_2_O_2_ production elevates intracellular ROS levels, leading to structural cellular damage.

### *In vivo* oxidation of hemoglobin by Spn*-*H_2_O_2_ selects for encapsulated pneumococci in the lung

Building on our *in vitro* observation that Spn-H_2_O_2_ oxidation of hemoglobin favors encapsulated pneumococci, we investigated the lung subcellular compartments where this selection occurs. Lung sections from mice infected with Spn were stained for capsule, DNA, and cell membranes, and analyzed by confocal microscopy. Notably, within the open central channel (lumen) of respiratory bronchioles in TIGR4-infected mice (Fig. 4A) or mice infected with the complemented TIGR4Ω*spxB-lctO* strain (Fig. 4C), nearly all pneumococci (98%) lacked a capsule (referred to as decapsulated or D-CPS) (Fig. 4D). Furthermore, D-CPS bacteria were the dominant population (88.4%) found inside the bronchiolar epithelium (Fig. 4A, 4C-D, and Supplementary Fig. 7). The proportion of CPS^+^ pneumococci increased significantly (by 42.1%) in Spn located within alveoli adjacent to bronchioles (Fig. 4D).

**Fig. 4.**
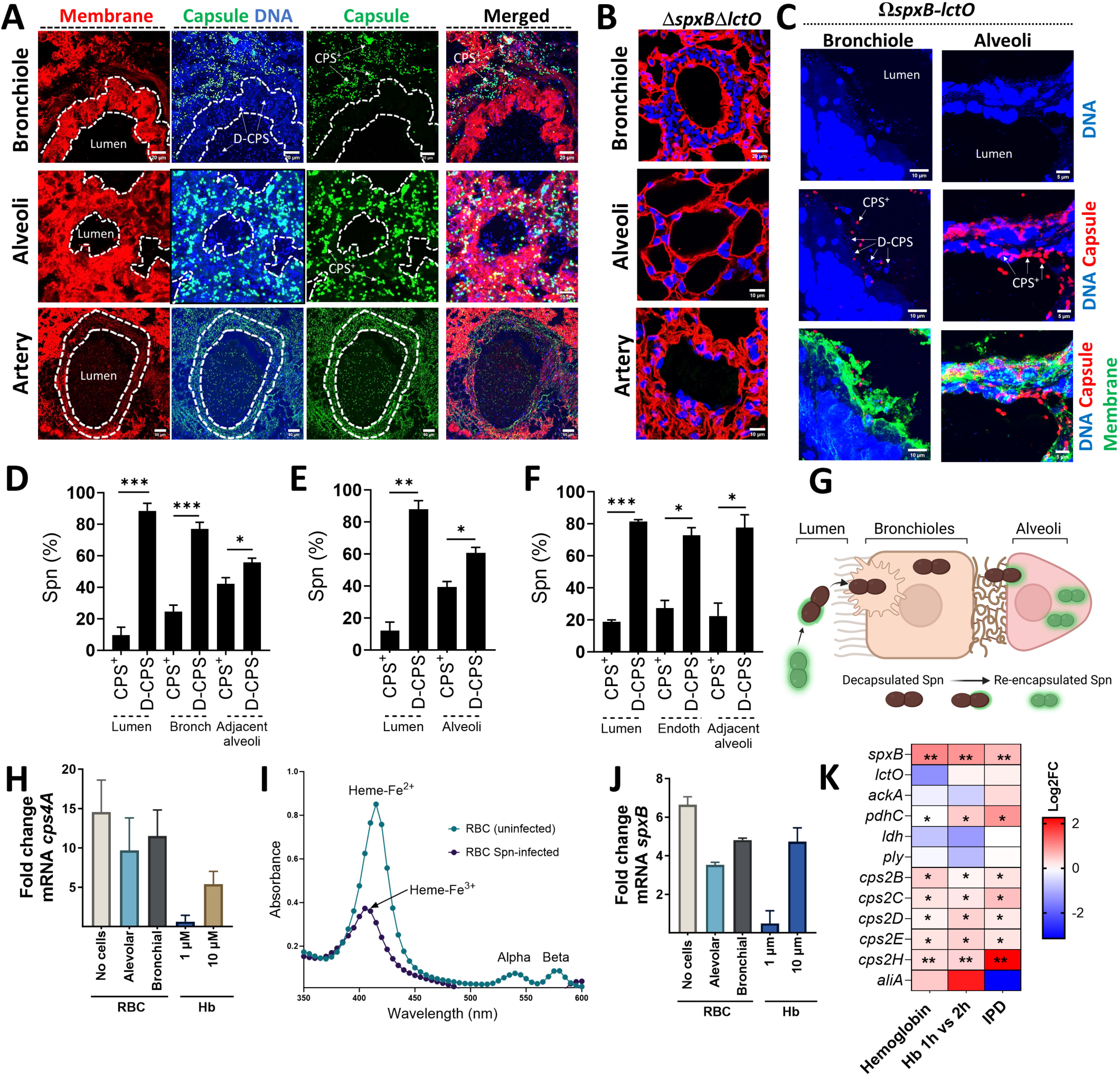
*S. pneumoniae* producing hydrogen peroxide encapsulates in the alveolar epithelium. Balb/c mice were intranasally infected with TIGR4 (∼10⁷ CFU) (A, D-F), TIGR4Δ*spxB*Δ*lctO* (B), or complemented TIGR4Ω*spxB*-*lctO* (C) and euthanized when moribund or at 96 h post-infection. Lung sections were stained for cell membrane, capsule, and DNA. Confocal microscopy was used to image bronchioles, alveoli, and pulmonary arteries and z-stack projections were obtained. Encapsulated (CPS^+^) and de-capsulated (D-CPS) pneumococci are indicated. A region of interest (ROI) was defined to delineate lumen and cellular compartments. DAPI and capsule fluorescence intensity, and CPS^+^or D-CPS pneumococcal population density (% Spn) were quantified in bronchioles (D), alveoli (E), and arteries (F) of TIGR4-infected mice. (G) Schematic depicting the hypothesized transition of CPS^+^ pneumococci from bronchioles to alveoli, involving de-capsulation (D-CPS) for bronchiolar invasion and subsequent re-encapsulation in the alveolar epithelium. (H-J) TIGR4 was cultured in THY (no cells), alveolar A549, or bronchial Calu-3 cells, with red blood cells (RBC), or in THY supplemented with hemoglobin (Hb). RNA was extracted, and *cps4A* (H) and *spxB* (J) gene transcription was analyzed. Hb release in THY-RBC cultures infected with TIGR4 (RBC Spn-infected) was assessed by spectroscopy (I), with indicated wavelengths for oxy-hemoglobin (Heme-Fe^2+^) and met-hemoglobin (Heme-Fe^3+^). (K) Heatmap visualization compares gene expression levels. Genes analyzed include those encoding pyruvate node enzymes, pneumolysin (*ply*) and capsule locus genes. Samples are from D39 cultures grown on THY-Hb^3+^ (Hemoglobin) vs. THY, THY-Hb^3+^ incubated for 2 h vs. 1 h and from *In vivo* TIGR4-infection model of invasive pneumococcal disease (IPD) compared with gene expression in the nasopharynx. Analysis was performed using the Mann Whitney test (D-F) and the non-parametric Wilcox test in (K) ns= not significant, *p<0.05, **p<0.01, ***p<0.001.

In stark contrast to the bronchiolar environment, the proportion of CPS^+^ pneumococci increased significantly (by 39.5%) within the alveoli, while the lumen (central channel) showed a marked decrease (Fig. 4A, 4C, and 4E). This trend continued as CPS^+^ were 27.3% when bacteria invaded the endothelial cells of pulmonary arteries (Fig. 4A and 4F). Notably, a similar pattern was observed in D39-infected mice, with these CPS^+^ bacteria concentrated on the bronchiolar membrane and in alveoli adjacent to bronchioles (Supplementary Fig. 5E). Comparable findings were noted in mice infected with serotype 3 strain WU-2 (data not shown). Since we began the experiment with a population of CPS^+^ pneumococci infected into the animals (Fig. 3B), these findings suggest that the bacteria lose their capsule (decapsulate) during bronchiolar invasion and then regain it (re-encapsulate) as they migrate deeper into the lung reaching the alveolar epithelium (Fig. 4G).

Motivated by the abundance of hemoglobin in the alveolar-capillary network, we investigated how its oxidation affects capsular gene expression. Incubating pneumococci (Spn) with red blood cells (RBCs), adding RBCs to alveolar cells, or bronchial cells all significantly increased transcription of *cps4A*, the first gene in the capsule locus, and *spxB* (Fig. 4H and 4J). Spectroscopic analysis of culture supernatants confirmed methemoglobin formation (heme-Fe^3+^) in Spn-infected RBCs-containing cultures, where most heme was already degraded (Fig. 4I). Importantly, only the higher concentration (10 µM) of hemoglobin mimicked the upregulation observed with RBCs (Fig. 4H and 4J), suggesting a dose-dependent effect. Previous studies reported similar findings, Spn grown in hemoglobin, mouse model of pneumonia but not in a carriage model, exhibited upregulated *spxB* and *cps* locus genes (Fig. 4K, Supplementary Fig. 6G). Notably, genes encoding pyruvate node enzymes (*lctO, ackA, ldh*) and the pneumolysin gene *ply* remained unaffected by hemoglobin exposure (Fig. 4K, Supplementary Fig. 6G). Similarly, ply expression remained unchanged in the H₂O₂-deficient TIGR4ΔspxBΔlctO mutant compared to wild-type TIGR4 under hemoglobin exposure^32^.

### Hemoglobin polymerization occurs in the lung during pneumococcal disease

We then investigated hemoglobin oxidation *in vivo*. Since hemoglobin oxidation can lead to its aggregation, we examined lung sections from Spn-infected animals for the presence of aggregated hemoglobin. We discovered aggregated hemoglobin, red blood cells (RBCs), CPS^+^ pneumococci, and bacteria associated with hemoglobin (Spn-Hb^+^) within alveoli (Fig. 5A, and Supplementary Fig. 8), the alveolar lumen (not shown), and the pulmonary artery (Fig. 5D). In contrast, mice infected with the H_2_O_2_-deficient TIGR4*ΔspxBΔlctO* strain did not exhibit aggregated hemoglobin or Spn-Hb^+^ bacteria in the alveoli (Fig. 5B) or pulmonary arteries (Fig. 5C). To assess the potential toxic effect of hemoglobin oxidation on Spn, we quantified the DNA area in both CPS^+^ pneumococci and Spn-Hb^+^ bacteria. The DNA area associated with Spn-Hb^+^ bacteria was significantly lower compared to that of CPS^+^ pneumococci (Fig. 5E), suggesting that hemoglobin oxidation may impair bacterial survival.

**Fig. 5.**
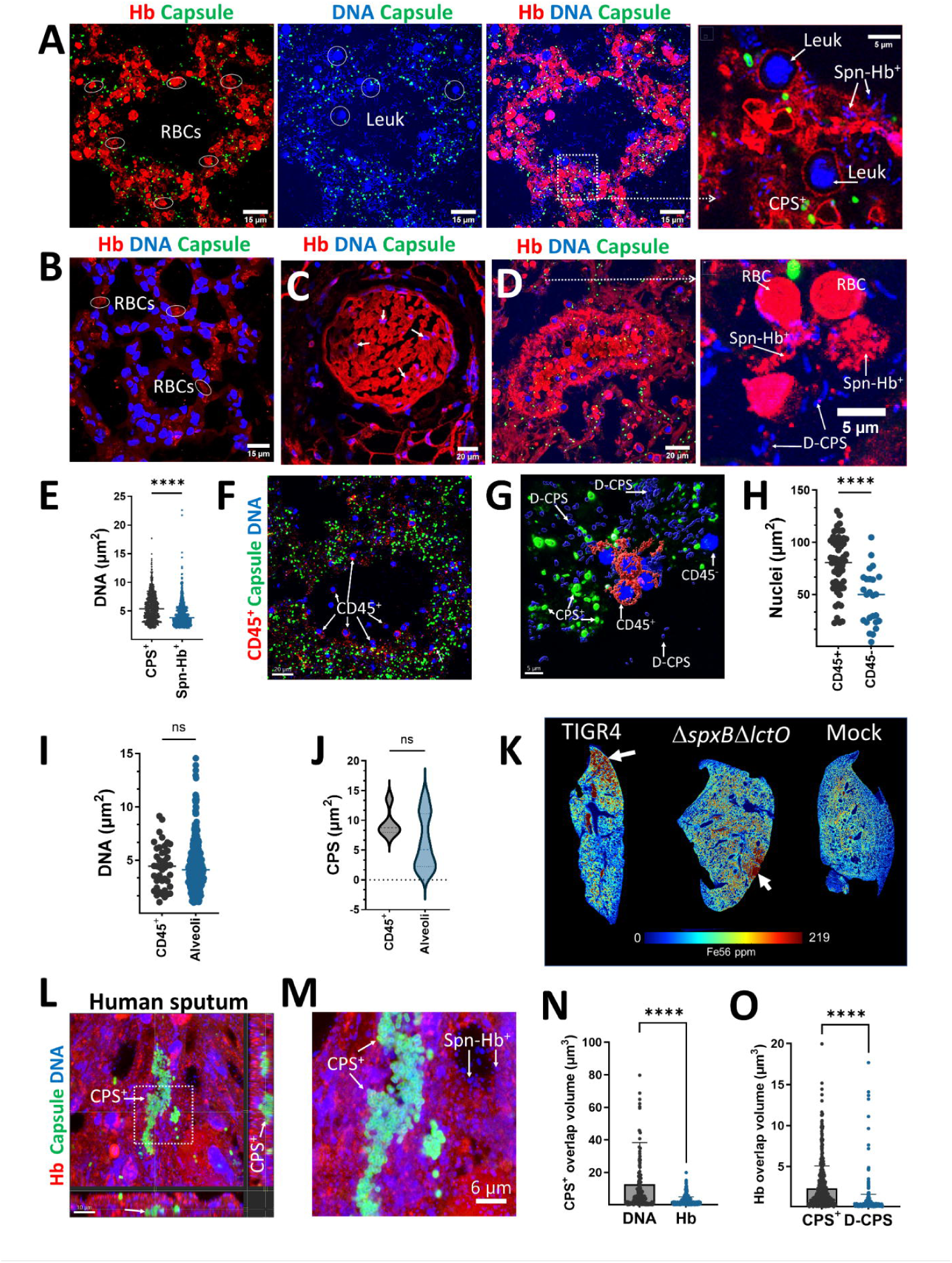
In vivo hemoglobin oxidation selects for encapsulated pneumococci. (A-D, F-H) Lung sections from Balb/c mice infected with TIGR4 or TIGR4Δ*spxB*Δ*lctO* were immunostained for hemoglobin, capsule, DNA and/or CD45^+^. Confocal microscopy z-stack projections of alveoli (A, B, and F) and pulmonary arteries (C and D) are shown. Red blood cells (RBC), leukocytes (Leuk), encapsulated (CPS^+^), de-capsulated (D-CPS) bacteria, hemoglobin-bound pneumococci (Spn-Hb^+^), and CD45^+^ leukocytes are indicated. (E) Quantification of DNA area within CPS^+^and Spn-Hb^+^ pneumococci in alveoli using Imaris software. Confocal micrographs (F) were used to generate (G) 3D masks for CD45^+^ and CD45^-^ cells, CPS^+^ and D-CPS pneumococci using Imaris software. (H) Nuclear area in CD45^+^ and CD45^-^ cells. (I) DNA area and (J) capsule area within CPS^+^ pneumococci, in close proximity (∼1 µm) to CD45^+^ cells, or farther and classified as alveolar pneumococci. At least three lungs from infected animals were analyzed. Micrographs and quantitative data are presented from a representative animal. (K) Elemental mapping of ^56^Fe^+^ in lung sections by LA-ICP-TOF-MS, with iron accumulation indicated by arrows. (L-O) Human sputum stained for hemoglobin, pneumococcal capsule and DNA. (L) Confocal optical XY middle section and corresponding XZ (underneath), and YZ (side) sections; (M) enlarged region. (N) Overlap volume analysis of the capsule and DNA and capsule and hemoglobin. (O) Overlap volume analysis of hemoglobin (Hb) and DNA from CPS+ and hemoglobin and DNA from non-encapsulated bacteria (D-CPS). Statistical significance was determined using the Mann Whitney test (E, G, I, J); ns= not significant, ****p<0.0001.

In alveoli with extensive hemoglobin polymerization, we observed nuclei resembling those of white blood cells (leukocytes), but no nuclei from other cell types such as type I and II pneumocytes or endothelial cells were present (Fig. 5A, Supplementary Fig. 8). To confirm that these were leukocytes migrating after tissue damage, lung sections were stained with an anti-CD45 antibody (Fig. 5F). This revealed that a majority of the cells (median, 56%) were CD45^+^, indicating immune cells. Compared to nuclear area of CD45^+^ cells, the nuclear area of these CD45^-^ cells was significantly shrunken, suggesting cytotoxicity (Fig. 5G).

We then examined pneumococci in close proximity, within ∼1 µm, to CD45^+^ cells (Fig. 5H). The area of both the DNA (Fig. 5I) and capsule (Fig. 5J) of CPS^+^ pneumococci near CD45^+^ cells was similar to those of bacteria farther away in the alveoli. This suggests that capsule expression was not directly influenced by the presence of immune cells.

Since hemoglobin oxidation in the alveoli leads to heme degradation and iron release^11, 33^, we expected elevated iron levels in the lungs. Using high-resolution laser ablation inductively coupled plasma time-of-flight mass spectrometry (LA-ICP-TOFMS), we quantified iron levels in lung sections from infected and uninfected mice. These analyses revealed high iron concentrations in animals infected with H_2_O_2_-producing pneumococci, with iron localized in areas exhibiting tissue damage (Fig. 5K, Supplementary Fig. 9). In contrast, mice infected with the H_2_O_2_-deficient TIGR4*ΔspxBΔlctO* strain only displayed increased iron levels in areas with abundant inflammatory infiltrate (Supplementary Fig. 9). These findings collectively demonstrate that *in vivo* hemoglobin oxidation triggers its polymerization, facilitates the formation of Spn-Hb^+^ pneumococci, and promotes bacterial encapsulation.

Next, hemoglobin oxidation was investigated in sputum from a pneumococcal pneumonia patient^34^. Ultrastructural analysis revealed aggregation of CPS^+^ pneumococci embedded in the sputum sample (Fig. 5L) and presence of Spn-Hb^+^ bacteria throughout (Fig. 5M). Hemoglobin signal was detected surrounding encapsulated bacteria, as quantified by overlap volume analysis. Due to bacterial embedding within the aggregate, hemoglobin-capsule interactions were significantly reduced compared to capsule-DNA interaction (Fig. 5N). Furthermore, hemoglobin overlap with DNA from Spn-Hb^+^ bacteria was significantly lower than that from CPS^+^ pneumococci (Fig. 5O), suggesting hemoglobin-induced toxicity in Spn-Hb^+^ bacteria.

### Both H_2_O_2_ and the capsule protect pneumococci from heme toxicity

Since heme exposure can kill a subpopulation of pneumococci, we investigated the mechanisms of heme resistance. Minimum bactericidal concentration (MBC) assays revealed that ≥10 µM heme was needed to kill 99.97% of wild-type pneumococci (Fig. 6A, Fig. 6E, Fig. 6G, and Supplementary Fig. 10A). Culture supernatants from pneumococci surviving exposure to lower heme concentrations (≤7.5 µM) showed significant heme degradation, while cultures exposed to a lethal dose (12.5 µM) retained the characteristic heme peak in spectral analysis (Fig. 6C and 6D). These findings suggest that pneumococci can degrade heme at low concentrations, potentially protecting themselves from its toxic effects. Labile heme levels in the bloodstream and lungs are ∼0.2 µM^35, 36, 37^. Therefore, H_2_O_2_ production by pneumococci might be particularly important for defense against heme toxicity at these physiological concentrations.

**Fig. 6.**
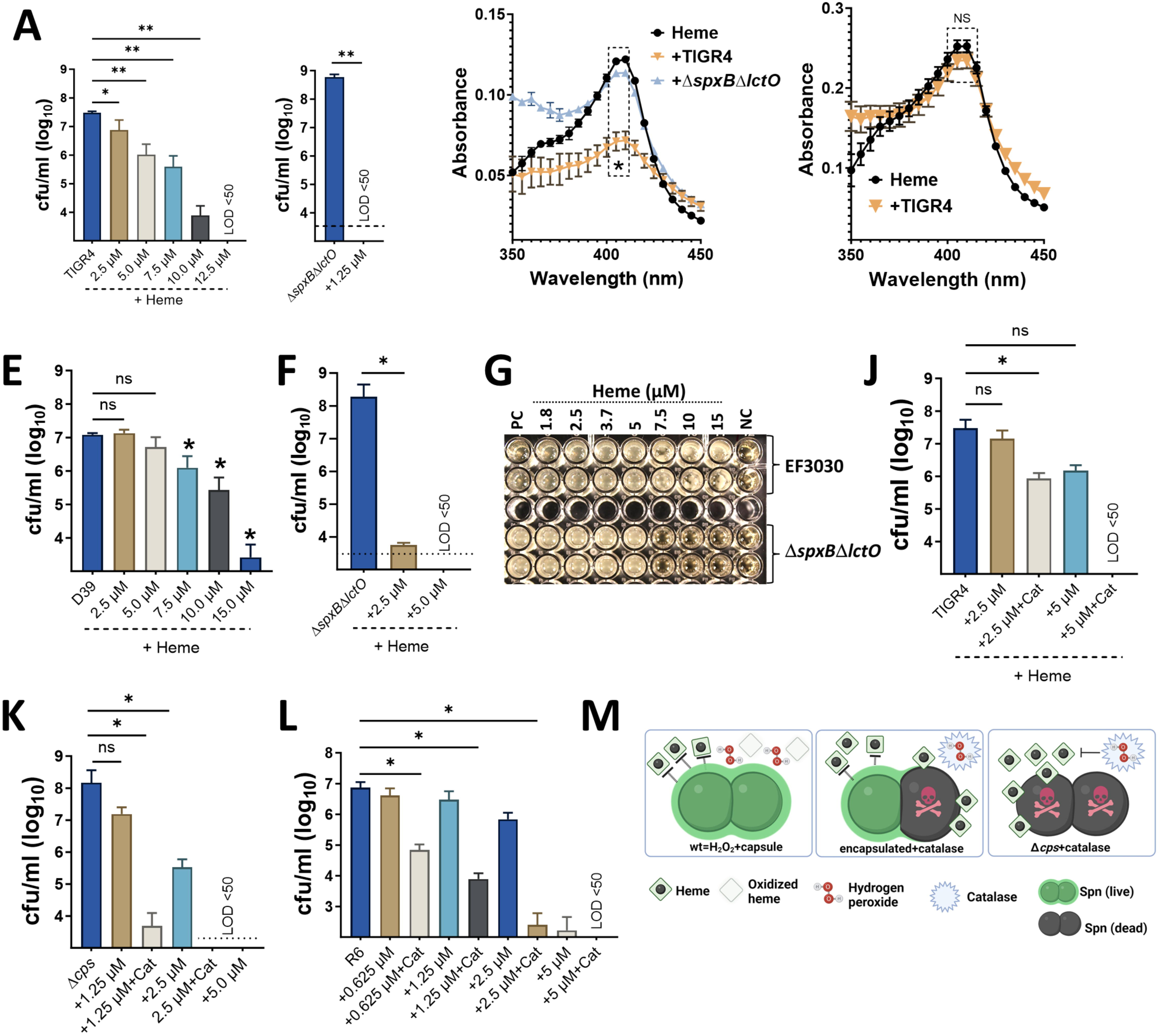
Hydrogen peroxide and the capsule protects pneumococci against heme toxicity. TIGR4 (A), TIGR4Δ*spxB*Δ*lctO* (B), D39 (E), or D39Δ*spxB*Δ*lctO* (F) were incubated with or without heme (4 h). Bacterial density was measured. Supernatants from heme challenge [(5 μM), C] or [(12.5 μM), D] were collected for spectral analysis. (G) EF3030 or EF3030Δ*spxB*Δ*lctO* were cultured in a 96-well plate with heme-supplemented THY and incubated overnight. NC: uninfected THY; PC: untreated THY with pneumococci. TIGR4 (J), TIGR4Δ*cps* (K), or R6 (L) were cultured in heme supplemented THY with or without catalase (400 U/mL). Cultures were incubated (4 h) and density measured by dilution and plating. (M) Proposed model for capsule and H_2_O_2_ protection against heme toxicity. Limit of detection (LOD) <50 CFU/mL. Data represent the mean ± SEM of two technical replicates from three independent experiments. Statistical analysis: Mann-Whitney U test was used to compare Soret peak absorbance (405 nm) between uninfected and pneumococci-infected media (B-D); (A, E, F, J-L) One-way ANOVA with Dunnett’s multiple comparison test. ns: not significant, *: p < 0.05, **: p < 0.01.

To directly test this hypothesis, we performed heme susceptibility assays in cultures lacking H_2_O_2_ production. H_2_O_2_-defective strains (*ΔspxBΔlctO*) were significantly more susceptible to heme, with an MBC below 2.5 µM (Fig. 6B, Fig. 6F and Supplementary Fig. 10B). Similarly, treating wild-type pneumococci with catalase, an enzyme that degrades H_2_O_2_, also rendered them susceptible to 2.5 µM heme (Fig. 6J). Spectral analysis confirmed the presence of intact heme in supernatants from cultures lacking H_2_O_2_ (Fig. 6C).

We then explored whether the pneumococcal capsule contributes to heme resistance, in addition to the established role of H_2_O_2_ production. Capsule-deficient strains (TIGR4Δ*cps* and R6) displayed increased heme susceptibility compared to wild-type strains (MBC of 5 µM for mutants vs. >10 µM for wild-type) despite producing normal H_2_O_2_ levels (Fig. 1B, 6K-L, and Supplementary Fig. 1A). Furthermore, scavenging H_2_O_2_ with catalase in these capsule-deficient strains dramatically reduced the MBC of heme to 1.25 µM or 0.625 µM, respectively (Fig. 6K and 6L).

To assess labile heme as a defense strategy, mice were infected with a heme-susceptible pneumococcal strain (TIGR4*ΔspxBΔlctO*) after depletion of complement factor 3 (C3) using cobra venom factor (CoVF). C3 depletion increases susceptibility to bloodstream infections^38^. We reasoned that if heme offered protection, CoVF-treated mice would still be resistant (Suppl. Fig. 4D). While some CoVF-treated mice lost weight (Suppl. Fig. 4E), likely due to CoVF itself, and as per protocol were euthanized and considered in the mortality group (Suppl. Fig. 4F), there were no significant differences in bacterial colonization or dissemination compared to controls (Suppl. Fig. 4G).

Collectively, these *in vivo* and *in vitro* experiments demonstrate that both the capsule and H_2_O_2_ production contribute to pneumococcal heme resistance (Fig. 6M). Loss of either factor increases heme susceptibility, and the absence of both renders pneumococci highly susceptible to heme toxicity at concentrations below 1.5 µM (Fig. 6M).

## Discussion

Spn thrives in the lung’s alveolar-capillary network, exploiting host oxygen, carbohydrates, and iron from heme-hemoglobin. However, this study demonstrates that Spn’s aerobic metabolism releases H_2_O_2_, unintentionally causing host damage. *In vitro*, Spn-H_2_O_2_ reacts with Fe^2+^, generating hydroxyl radicals (^•^OH^-^)^39, 40^ and oxidizes hemoproteins (hemoglobin, cytochrome C) to labile heme and tyrosyl radicals^9, 11, 12^. Our findings reveal these Spn-H_2_O_2_ driven oxidative reactions as a major contributor to lung toxicity. Additionally, neutrophil infiltration in response to pneumococcal infection^41^ produces hydrogen peroxide, potentially synergizing with Spn-H₂O₂ to exacerbate oxidation of heme-containing proteins within the alveolar-capillary microenvironment. Hemoglobin oxidation triggers cytosolic protein degradation and facilitates Spn differentiation into encapsulated bacteria, primarily within alveoli and pulmonary artery endothelial cells. These mechanisms represent novel targets for intervention against pneumococcal disease, which claims millions of lives annually.

Spn-H_2_O_2_ oxidation of hemoglobin induces globin chain aggregation (α and β) under physiological infection conditions, potentially aiding pneumococcal suspension in lung tissue^22, 23^. This contrasts with *in vitro* studies reporting preferential crosslinking of either α or β globin by H_2_O_2_. The oxidation also generates methionine sulfoxide (MetOH) in the β chain, potentially contributing to globin collapse^23^. Free hemoglobin is normally bound by haptoglobin (Hp) for macrophage uptake via CD163 receptors. However, oxidized hemoglobin exhibits reduced Hp binding, weakening its interaction with macrophages^22, 42^. In our mouse model, oxidized hemoglobin bound to a subset of non-encapsulated Spn-Hb^+^ bacteria, while the majority localized near PMNs. This suggests membrane-bound hemoglobin in Spn-Hb^+^ bacteria trigger an immune response, supported by the abundance of CD45^+^ leukocytes surrounding Spn-Hb^+^ in severe pneumonia.

Pneumolysin (Ply), a pneumococcal toxin, may not fully explain the complex lung pathology observed in pneumonia^43, 44, 45, 46, 47, 48^. Spn-H_2_O_2_ shows promise as a mediator of both cell death and pneumococcal disease in animal models^28, 45, 49, 50, 51, 52^. Notably, the co-activation of apoptosis and pyroptosis markers suggests a multifaceted cytotoxicity mechanism^53, 54, 55, 56^. Our study identified a loss of cellular structure in mouse lung regions exhibiting severe alveolar and bronchial cytotoxicity, harboring intracellular pneumococci. Notably, epithelial cell nuclear DNA was absent, while infiltrated CD45^+^ cells retained nuclei. *In vivo* cytotoxicity was partially attributed to Spn-H_2_O_2_ produced by pneumococci in the lungs. Furthermore, Spn-H_2_O_2_ was delivered intracellularly to cultured human alveolar cells. While Spn-H_2_O_2_ can directly oxidize proteins, LA-ICP-TOFMS analysis revealed elevated iron levels in these cytotoxic regions. This suggests a potential role for the Fenton reaction in generating hydroxyl radicals (^•^OH^-^) and subsequent DNA damage, contributing to the observed toxicity^40, 57^.

Our study unveils a mechanism linking H_2_O_2_-producing pneumococci to lung cell death and structural collapse. Infection of human alveolar or bronchial cells with these bacteria triggered the degradation of key cytoskeletal proteins (β-tubulin, β-actin) and cytosolic GAPDH (glycolysis) ^12, 31^. This correlated with microtubule collapse and cell swelling, potentially mimicking damage observed in cardiomyocytes exposed to high H_2_O_2_^30^. Mechanistically, Spn-H_2_O_2_ likely targets cysteine residues in α and β-globin chains, weakening tubulin-tubulin interactions and inhibiting polymerization^30, 58,59^. Similarly, H_2_O_2_ can directly oxidize the catalytic cysteine (Cys152) of GAPDH, leading to S-glutathionylation and subsequent cell death^31, 60, 61^. This process holds therapeutic potential, as studies suggest pharmacological induction of GAPDH oxidation shows promise for treating tumors^62^. Furthermore, antioxidant treatment in animal models and human trials improves survival rates from pneumococcal infections^63, 64, 65^. Spn-H_2_O_2_ oxidation of hemoglobin triggers a cascade of detrimental events within the lung. This process releases heme, leading to its degradation and the formation of highly reactive tyrosyl radicals^12, 17, 18, 66, 67^. Heme destruction is well-documented for its toxicity. For instance, the accumulation of labile heme during hemolytic diseases like malaria and sickle cell disease (SCD) is linked to inflammation, blood clot formation (prothrombotic complications), and a form of cell death called PANoptosis^15, 68, 69^. Importantly, the oxidation of other heme-containing proteins like myoglobin, cytochrome C (Cytc), and methemoglobin (Hb-Fe3+) also generates these toxic tyrosyl radicals^12, 70, 71, 72, 73^. Collectively, our findings unveil novel intervention strategies. By addressing oxidative reactions in pneumococcal pneumonia or by stimulating the natural antioxidant response of lung cells through pharmacological approaches, we may unlock promising therapeutic possibilities.

## Material and Methods

The Materials and Methods section is provided in supplementary materials. It can be accessed online for further details.

## Acknowledgements

This study was supported in part by grants from the National Institutes of Health including R21AI144571, 1R01AI175461-01A1 (JEV). BA was supported by a Fulbright scholarship awarded by the US Department of State. Studies of confocal microscopy, histology, and Oroboros O2k FluoRespirometry, were supported by grants from the National Institute of General Medical Sciences (NIGMS) of the National Institutes of Health under Award Numbers P20GM121334 and P20GM104357. JEV is also supported by a grant from NIGMS through the Molecular Center of Health and Disease (1P20GM144041-01A1 7651). Laser ablation elemental imaging was performed at the Dartmouth Biomedical National Elemental Imaging Resource (BNEIR) supported by NIGMS R24GM141194. RNA-seq data analysis was performed through the UMMC Molecular and Genomics Facility is supported, in part, by funds from the NIGMS, including the Molecular Center of Health and Disease-COBRE (P20GM144041), Mississippi INBRE (P20GM103476) and Obesity, Cardiorenal and Metabolic Diseases-COBRE (P30GM149404) and Mississippi Center of Perinatal Research (1P20GM121334). The content is solely the responsibility of the authors and does not necessarily represent the official view of the NIH or the US Department of State. We thank Drs. Poonam Sharma, William Daley, and Timothy Allen from the Department of Pathology at the University of Mississippi Medical Center (UMMC) for providing the human sputum sample, and Dr. Allen for assistance with pathology scoring. We also appreciate the assistance of Dr. Antonino Baez (UMMC) for CPS purification and Jorge M. Vidal (UMMC) for conducting experiments with serotype 3 strains.

