## Supplemental information for "Heme-Mediated Selection of Encapsulated *Streptococcus pneumoniae* in the Lungs by Oxidative Stress"

**Supplementary:**

### RESOURCE AVAILABILITY

### Material and Methods

**Ethics Statement.** All experiments involving animals were performed with prior approval of and in accordance with protocol 1584 which was reviewed and approved by the UMMC Animal Care Committee. UMMC laboratory animal facilities have been fully accredited by the American Association for Accreditation of Laboratory Animal Care. Procedures were performed according to the institutional policies, Animal Welfare Act, NIH guidelines, and American Veterinary Medical Association guidelines on euthanasia.

The male and female 4-5-week-old Balb/c mice were obtained from Charles River Laboratories, Inc and allowed to acclimate for one week before the experimental challenge. At all times mice received food and water ad libitum.

**Bacterial strains and growth conditions.** All *Streptococcus pneumoniae* (Spn) strains used in this study are listed in the Table S2. Spn was cultured on BAP (Fisher Scientific) or BAP with 10 µg/ml gentamicin (Sigma-Aldrich) at 37°C for 24 h in a 5% $CO_2$ atmosphere. Colonies were inoculated into Todd-Hewitt-broth with 0.5% yeast extract (THY) (BD) and grown until exponential phase ($OD_{600}$ 0.3, ~1 X $10^8$ CFU/ ml). THY broth was utilized in all the experiments. Isogenic mutants were cultivated using appropriate antibiotics.

**Cell culture lines.** Low passage, A549 human type II alveolar epithelial cells (ATCC, CCL-185), and MLE-12 mouse lung type II epithelial cells (ATCC, CRL-2110), and Calu-3 Cells (ATCC, HTB-55) were purchased from ATCC (American Type Culture Collection). The A549 and MLE-12 cells were cultured in Dulbecco's M0dified Eagle's Medium (DMEM) (Corning) media with 2 mM L-glutamine (Gibco), 10% fetal bovine serum (FBS) (R&D systems, Bio-techne), 100 units/ml penicillin/streptomycin solution (Gibco). Calu-3 cells were cultured in Eagle’s Minimum Essential Medium (American Type Culture Collection, ATCC) supplemented with 10% FBS and 100 U/mL of penicillin-streptomycin (Gibco). Cell passages were performed by trypsinization with Trypsin-EDTA (0.25%), phenol red (Gibco) and seeded in the experiment specific device. All experiments described hereafter were performed with cells that had been grown for 8-10 days in the specified device.

**Preparation of inoculum for *in vitro* and for animal experiments.** Inoculum for the *in vitro* studies was prepared essentially as previously described[1, 2, 3]. Briefly, an overnight BAP culture of the Spn strain was used to prepare a bacterial suspension in sterile phosphate buffered saline [(PBS), pH=7.4] (Fisher BioReagents). This fresh bacterial suspension was inoculated to a final $OD_{600}$ of ~0.1 and the suspension contained ~5.15x$10^8$ cfu/ml. Aliquots of these suspensions were routinely diluted and plated to confirm bacterial counts (cfu/ml). Unless otherwise noted, all *in vitro* experiments described below were inoculated with fresh inocula, i.e., prepared right before the start of the experiment.

To prepare inoculant for mice studies, the Spn strain was inoculated in THY broth supplemented with 20% fetal bovine serum (FBS) (R&D systems, Bio-techne); this culture was grown for 4 h (i.e., early log phase) at 37°C in a 5% $CO_2$ atmosphere; the growth was monitored using a spectrophotometer Bio-Rad SmartSpec Plus. After this incubation time, the culture was centrifuged at 4500 g for 5 min and the pellet was resuspended in 10 ml PBS and washed once. The washed bacterial pellet was resuspended in PBS containing 10% (v/v) glycerol and stored at -80ºC until it was used. Aliquots of inocula were thawed from each batch to obtain the density by dilution and platting. All inocula were used within two months.

**Confocal microscopy studies.** All micrographs obtained in these experiments and throughout this study were obtained with a Nikon C2 laser scanning confocal microscopy system.

*(A) Studies of pneumococcal biofilms, and planktonic pneumococci, incubated with hemoglobin.* Pneumococci were inoculated into 8-well glass slides (Thermo Fisher Scientific, Nunc Lab-Tek II CC2) that had been added with THY broth supplemented with 10 µM hemoglobin (Sigma-Aldrich) and incubated at 37°C in a 5% $CO_2$ atmosphere. At the end of the incubation time, planktonic pneumococci were removed, washed two times with PBS, and fixed with 4% paraformaldehyde (PFA) (Sigma-Aldrich) for 15 min at room temperature. Biofilms were washed two times with PBS and then fixed with PFA as above. Planktonic pneumococci, or biofilms, were then washed two times with PBS and with 2% bovine serum albumin (BSA) (Roche) for 30 min at room temperature. Planktonic bacteria were stained in suspension whereas biofilms were stained on the 8-well glass slides. Pneumococci were then stained for 1 h at room temperature with serotype-specific polyclonal antibody (Statens Serum Institute, Denmark) (40 µg/ml) that had been previously labeled with Alexa-488 (anti-S4-A488) (Thermo Scientific) or Alexa-555 (anti-S2-A555, or anti-S19-A555) (Molecular Probes) and with a mouse monoclonal anti-hemoglobin antibody labeled with Alexa-546 [(anti-Hb-A546), 20 µg/ml] (Santa Cruz Biotechnology). Stained planktonic bacteria or biofilms were finally washed three times with PBS. Planktonic pneumococci

dropped on microscope slides and stained biofilms were air dried and then mounted with ProLong Diamond Antifade mounting medium containing DAPI (Molecular Probes).

*(B) Confocal microscopy studies of lung tissue.* Deparaffinized sections of lung tissue ~5 µm mounted on microscope slides were washed three times with PBS and the preparations were blocked with 2% bovine serum albumin (BSA) for 1 h at room temperature. These preparations were then incubated for 1 h with serotype-specific polyclonal antibody (Statens Serum Institute, Denmark) (40 µg/ml) that had been previously labeled with Alexa-488 (anti-S4-A488) or Alexa-555 (anti-S2-A555) (Molecular Probes) and wheat germ agglutinin conjugated to Alexa-488 or Alexa-555 [(WGA), 5 µg/ml]. Some lung tissue specimens were stained with an anti-S4-A488 and a mouse monoclonal anti-hemoglobin antibody labeled with Alexa-546 [(anti-Hb-A546), 20 µg/ml] (Santa Cruz Biotechnology) for 1 h at room temperature. Stained tissues were finally washed three times with PBS, air dried and then mounted with ProLong Diamond Antifade mounting medium containing DAPI (Molecular Probes).

*(C) Studies of pneumococci incubated with human alveolar cells in the presence of hemoglobin.* Human A549 cells grown on an 8-well chamber (Thermo Fisher Scientific, Nunc Lab-Tek II CC2) were washed three times with PBS and then added with infection medium that had been supplemented with 10 µM hemoglobin. Pneumococci were infected and incubated at 37°C in a 5% $CO_2$ atmosphere. At the end of the incubation time, planktonic pneumococci that had been removed, and biofilms were washed, fixed with PFA and the capsule of pneumococci, DNA, and hemoglobin were stained essentially as detailed above.

(D) *Confocal studies of human sputum.* Human sputum from a de-identified patient with microbiologically and radiologically confirmed pneumococcal pneumonia was utilized for confocal microscopy. IRB approval was obtained for the use of this de-identified specimen. The sample was collected according to institutional guidelines by the University of Mississippi Medical Center's clinical laboratory and stored at -80°C until processed. Spn was confirmed in the specimen by *lytA*-based real-time PCR, as previously described[4, 5], with a quantification of $3.95\times10^4$ genome equivalents/ml. The sputum was stained for 1 hour with an Alexa-488-conjugated polyclonal anti-Spn antibody (Omni; Statens Serum Institute, Denmark; 40 µg/mL) and an Alexa-546-conjugated mouse monoclonal anti-hemoglobin antibody (Santa Cruz Biotechnology; 20 µg/mL). Samples were air-dried and mounted with ProLong Diamond Antifade Mountant with DAPI (Molecular Probes). This human sputum sample was part of the project “Mechanistic Studies of Broadly Reactive Antibodies for Pneumococcal Infection,” reviewed by the University of Mississippi Medical Center IRB (FWA#00003630) and deemed exempt from formal review. The IRB determined it meets exemption criteria under 45 CFR 46.102(l) as it does not involve research with human subjects, requiring no further ethical review.

**Quantification of pneumococci expressing capsule.** Aliquots of inocula prepared for animal studies, as detailed above, were thawed in ice and stained with a serotype-specific polyclonal antibody (40 µg/ml) for 1 h at room temperature. Stained pneumococci were pelleted down by centrifugation at 21,130 x g for 5 min, washed two times with PBS and then fixed with 4% PFA that was incubated for 15 min at room temperature. Fixed bacteria were washed again two times with PBS, resuspended in PBS and air dried on a microscope slide. Preparations were mounted with ProLong Diamond Antifade mounting medium containing DAPI and micrographs were obtained by a confocal microscope. Capsule was quantified from confocal micrographs using ImageJ (NIH). The quantification process consisted of separating the DAPI channel (bacterial DNA) and the green channel (encapsulated pneumococci), quantifying the bacteria in the DAPI channel using the 'find maxima' function, then overlaying the green channel against the DAPI channel using the 'image calculator' function to confirm encapsulation before quantifying the green channel in the same manner. The 'prominence' setting was adjusted to filter out redundant counts, and the output was selected as 'single points'**.** Parameters were adjusted to reduce background noise. Absolute number of bacteria (encapsulated Spn/Spn) were obtained and utilized to calculate the percentage of pneumococci that expressed capsule by using Microscopy Image Analysis Software – Imaris (Oxford, UK).

**Adhesion assay.** Adhesion of pneumococci to A549 cells and MLE-12 cells was assessed as follows: Cells grown for 8-10 days on a 6-well microplate were washed three times with sterile PBS to remove the antibiotic and added with 1 ml of infection medium, which consisted of DMEM supplemented with 2 mM L-glutamine, HEPES buffer (100 mM) and 5% fetal bovine serum (FBS). Cells were infected with pneumococci and incubated for the indicated time points, after which infected cells were washed two times with PBS to remove unbound bacteria. Cells and pneumococci were detached by sonication (Branson Ultrasonic 2800) for 30 s and after homogenization in PBS, the suspensions were diluted and plated to obtain bacterial accounts.

**Purification of the soluble fraction of human alveolar cells infected with pneumococci.** A549 cells were infected as above and incubated for 4, 8, 12, 16 or 24 h at 37°C in 5% $CO_2$ atmosphere. Uninfected cells were used as a control. After incubation, cells were gently scrape off the 6-well microplate and washed twice with ice-cold PBS by centrifugation cycles of 300 x g for 5 min at 4°C. Washed cells were lysed with RIPA buffer supplemented with 1X complete protease inhibitor

(Roche) incubated 30 min on ice. After incubation, lysed cells were centrifuged at 13,500 x g for 20 min at 4°C and the soluble fraction was collected. The protein concentration of the soluble fraction was determined with the BCA protein assays (Thermo Fisher Scientific) as recommended by the manufacturer.

**Quantification of intracellular reactive oxygen species.** To quantify reactive oxygen species (ROS) in infected cells the CellROX Green Reagent from Invitrogen was used. Human alveolar A549 cells were seeded in an 8 well chamber slide, and once confluent, cells were infected with Spn or mutant derivatives and incubated for 6 h at 37°C in a 5% $CO_2$ atmosphere. After incubation, planktonic pneumococci were removed and infected cells were incubated for 30 min with fresh infection medium containing the CellROX Green Reagent at a concentration of 5 μM. Cells were then washed three times with PBS and fixed with 2% PFA for 15 min after which the cells were washed two times with PBS, then air-dried, and mounted with ProLong Diamond Antifade mountant containing DAPI (50 μg/ml). Cells were immediately imaged by confocal microscopy. Quantification of intracellular ROS from confocal micrographs was performed using ImageJ version 1.3t (National Institutes of Health, NIH) and done by taking intensity spectrum measurements of a region of interest in the green channel (GFP) and obtaining area under the curve. Values were expressed by creating a ratio of ROS (GFP): DAPI intensity within the same samples.

**Bacterial cell fractionation.** To investigate heme or hemoglobin uptake, the subcellular fractions of TIGR4, TIGR4*ΔspxBΔlctO*, and TIGR4Δ*glnP* without or with hemoglobin, heme growth conditions were prepared by the method as previously described[6, 7] with minor modification. Briefly, grown bacterial cells were pelleted by centrifugation at 7,500 x g for 10 min at room temperature. The supernatant was discarded, cells were washed once with ice-cold PBS to remove cell debris, and resuspended in 1 ml protoplast buffer (1X protease inhibitor cocktail [Roche], 200 U/µl mutanolysin, 2 mg/ml lysozyme in a 20% sucrose-10 mM Tris [pH 7.5]- 50 mM MgCl2 buffer) and incubated at 37°C for 2 h with shaking. The bacterial endogenous autolytic enzymes hydrolyze the peptidoglycan layer of Spn activated by MgCl2, and the sucrose maintains the protoplasts intact. After incubation, the suspension was centrifuged at 21,130 x g for 10 min at 4°C, and the supernatant containing cell wall extract fraction was collected and kept at -80°C. The pellet protoplasts were resuspended in a 20% sucrose-10 mM Tris [pH 7.5]- 50 mM $MgCl_2$ buffer supplemented with 1X protease inhibitor cocktail [Roche] and lysed by sonication. The lysed cell suspension was centrifuged at 6000 x *g* for 10 min at 4°C to remove any particulate material. The supernatant-contained cytoplasmatic membrane and cytoplasmatic proteins were separated by ultracentrifugation at 25,000 x g for 3 h at 4°C. After centrifugation, the supernatant was kept as cytoplasmatic fraction, and the pellet cytoplasmatic membrane fraction resuspended in TN buffer and stored for further processing.

**Western blotting.** Equivalent amounts of cellular or subcellular fractions, as specified in the appropriate section, from bacteria, infected cells, or lung specimens were combined with 4x reducing sample buffer and boiled for 5 min at 100°C. The mixture was loaded into a 12% Mini-Protean TGX precast gels (Bio-Rad), which were run for 2 h at 90 V. Gels were transferred to nitrocellulose membranes using the recommended transfer buffer supplemented with 20% ethanol and the Trans-Blot Turbo transfer system. Transferred membranes were blocked for 1 h at room temperature in a solution of Tris-buffered saline supplemented with 0.1% Tween 20 (TBST) and 5% non-fat dry milk. The membrane was then incubated overnight at 4°C with a primary antibody diluted in TBST containing 5% non-fat dry milk. The following primary antibodies were used: anti-β tubulin at 1.6 µg/µl (Proteintech, 10068-1-AP), anti-β actin at 1.6 µg/µl (Proteintech, 66009-1-Ig), anti-GAPDH at 2 µg/µl (Proteintech, 60004-1-Ig), anti-catalase at 1.2 µg/µl (Proteintech, 21260-1-AP), anti-hemoglobin α chain mAb at 2 µg/µl, anti-hemoglobin β chain mAb at 2 µg/µl or an anti-hemoglobin polyclonal antibody at 10 µg/µl (Invitrogen).

The next day, the membrane was washed three times for 5 min each with TBST, in a rocking platform, and incubated for 1 h at room temperature with Starbright Blue 520 (Bio-Rad) anti-mouse, or anti-rabbit; this secondary antibody was diluted 1:2500 in 5% non-fat dry milk in TBST. The membrane was then washed six times for 5 min each with TBST. Blots were then imaged on a ChemiDoc MP imager (Bio-Rad) using Image Lab 5.0 software, with automatic exposure settings for chemiluminescence and high specificity, optimizing for bright bands. Densitometric analysis of the bands was performed with ImageLab software (Bio-Rad), normalizing to a selected target protein.

**Bacterial growth curves and studies of the minimum inhibitory concentration (MIC).** Bacteria were grown in 24-well plates poured with 1 ml fresh THY (with or without supplements) at 37°C in a 5% $CO_2$ atmosphere. The medium was then inoculated with fresh Spn inoculum, and the $OD_{600}$ was monitored every 20 minutes for 24 h using a BioTek Synergy H1 Multimode Reader (Agilent Technologies, Inc). Uninfected THY was always used as the blank and negative control. To assess the inhibition of growth by heme, a 2 mM stock solution of heme prepared in dimethyl sulfoxide (DMSO), was diluted in THY broth. This stock solution was used same day. Non-heme supplemented THY broth was used as a control. Pneumococci were inoculated and the $OD_{600}$ was monitored as above. Two technical replicates and three biological replicates were performed in these experiments.

The minimum inhibitory concentration (MIC) of heme was determined using a slightly modified broth microdilution method, as recommended by the Clinical Laboratory Standards Institute (CLSI)[8]. Briefly, a 2 mM stock solution of heme was diluted in THY broth and added in 96-well plates; note that we cannot use lysed horse blood, as recommended by the CLSI, because the lysate contains hemoglobin and heme. The 96-well plates were inoculated with pneumococci at ~10 x $10^8$ cfu/ml and incubated at 37°C in a 5% $CO_2$ atmosphere for 24 h. The MIC was recorded as the heme concentration of the well with no visible growth. MIC assays were performed with technical triplicates and two biological replicates.

To assess the viability of pneumococci challenged with heme, Spn was inoculated in 6-well plates that had been added with THY, as a control, or THY supplemented with heme. The samples were incubated for 2, 4, or 6 h at 37°C in a 5% $CO_2$ atmosphere. After incubation, bacteria were collected, diluted, and plated on BAP to obtain the density (cfu).

**Spectroscopic analysis of heme, or hemoglobin, when incubated with pneumococci.** Spn was inoculated in THY broth or THY supplemented with heme (0.62, 1.25, 2.5, 5, 10, or 20 μM) or hemoglobin (10 μM), and incubated for the indicated time at 37°C in a 5% $CO_2$ atmosphere. The culture supernatants were collected, and then centrifuged at 4°C for 5 min at 21,130 x g and transferred to a 96-well plate and analyzed by spectroscopy spanning 200 nm through 1000 nm using an BMG LabTech FLUOstar Omega spectrophotometer.

**Qualitative assessment and quantification of hydrogen peroxide.** Spn hydrogen peroxide production was determined with qualitative and quantitate methods. Qualitative detection of $H_2O_2$ production by Spn was performed as previously described[3, 9] using the Prussian blue agar assay. Shortly, Spn strains or culture supernatants (10 μl), obtained as described above, were inoculated or spotted, respectively, onto Prussian blue agar plates and these plates were incubated at 37°C in a 5% $CO_2$ and ~20% $O_2$ atmosphere overnight (Spn) or for 4 h (supernatants). The plates were then photographed with a Canon Rebel EOS T5 camera system, and digital pictures were analyzed.

$H_2O_2$ was quantified from supernatants of Spn cultures that had been infected in THY (with or without supplements) and incubated at 37°C in a 5% $CO_2$ and ~20% $O_2$ atmosphere for the indicated time. At the end of the incubation, culture supernatants were collected and centrifuged at 4°C for 5 min at 21,130 x g, and then transferred to new tubes that were placed on ice. Some supernatants were filter sterilized using a 0.22-μm syringe filter (Genesee Scientific) and kept on ice. Collected supernatants were processed within 30 min by dilution with 1X reaction buffer from the Amplex Red $H_2O_2$ assay kit (Molecular Probes), and $H_2O_2$ levels were quantified according to the manufacturer's instructions using an BMG LabTech FLUOstar Omega spectrophotometer.

$H_2O_2$ was quantified from lungs removed 24 h post-infection with pneumococci or mock-infected. Lungs were homogenized within 20 minutes after removal using a LabGen Series 7 Homogenizer (Cole-Parmer) and $H_2O_2$ was quantified within 30 min. $H_2O_2$ concentration was measured using the Amplex Red (10 μM) which is converted to resorufin (highly fluorescent) by the reaction of $H_2O_2$ and horseradish peroxidase (HRP, 1U/ml). Superoxide dismutase (SOD, 5U/ml) was also added to convert superoxide (SO) to $H_2O_2$. The rate of $H_2O_2$ production was obtained by calibrating the fluorescence response of known concentrations of $H_2O_2$ in the Oroboros O2k FluoRespirometer. Additionally, the end point absorbance was measured using a SpectraMax M5 plate reader at 570 nm with a separate calibration curve for the absorbance.

**In-gel heme detection assay.** Bacterial cytosolic or membrane fractions, or supernatants harvested from experiments incubating pneumococci with hemoglobin were combined with nonreducing loading buffer. Samples were not heated. The mixtures were loaded into a nondenaturing 12% Mini-Protean TGX precast gel (Bio-Rad), which was run for 2 h at 100 V in running buffer lacking sodium dodecyl sulfate (SDS). The gel was then stained using a published protocol[10]. Briefly, after electrophoresis, gels were immersed in a methanol-sodium acetate solution (pH 5) and incubated at room temperature on a rocking platform (VWR) at a speed of 1 for 2 min. After incubation, a solution of o-dianisidine-sodium acetate (pH 5) was added to the gel, which was incubated for 20 min in the dark. To visualize heme in the gel, 3% hydrogen peroxide was added, and the reaction was stopped by washing with distilled water. The gel was then photographed with a Canon Rebel EOS T5 camera system, and digital pictures analyzed.

**Quantification of hemoglobin and heme.** The concentration of heme or hemoglobin, was determined by using the QuantiChrom™ Heme Assay Kit or the QuantiChrom™ Hemoglobin Assay Kit (BioAssay), respectively. Heme and Hemoglobin levels were determined using manufacturer's instructions, calibration standards provided in each kit and using a BMG LabTech FLUOstar Omega spectrophotometer.

**Purification of Standard Capsular Polysaccharide (CPS).** CPS was purified from Spn as previously described[1]. TIGR4 and D39 were cultured in Todd Hewitt broth with 0.5% yeast extract (THY) to an optical density at 600 nm ($OD_{600}$) of 0.5. Bacteria were harvested by centrifugation, and the pellets were resuspended in 50 mM citrate buffer containing 0.1% Triton X-100. After incubation at 42°C for 30 min, cell debris was removed by centrifugation. The CPS-containing supernatants were filtered through a 0.2-μm Nalgene Rapid-Flow filter. To remove nucleic acids, the filtrates were mixed with one-fourth volume of ethanol and stirred at 4 °C for 2 h. CPS was then precipitated by adding 80% ethanol and incubating at 4 °C for another 2 h. The precipitated CPS was collected by centrifugation, air-dried, and resuspended in sterile water. CPS concentration was determined using the phenol–sulfuric acid method[2]. Briefly, 25 μL of D-glucose standards or CPS samples were incubated with 125 μL of concentrated sulfuric acid at 4 °C for 15 min. Then, 25 μL of 5% phenol was added, and the mixtures were transferred to 96-well microplates. After shaking for 5 min and incubation at room temperature for 60 min, absorbance was measured at 490 nm.

**Quantification of Pneumococcal CPS by Enzyme-Linked Immunosorbent Assay (ELISA).** Pneumococcal CPS was quantified by indirect ELISA, as previously described[3],[4], with minor modifications. Bacteria were cultured in THY broth for 6 h and then heat-killed at 60 °C for 20 min. Bacterial densities were quantified under the same conditions to normalize CPS content across samples. Samples and purified CPS standards were diluted in carbonate-bicarbonate coating buffer (0.015 M $Na_2CO_3$, 0.035 M $NaHCO_3$) and added to slightly hydrophilic, flat-bottom 96-well plates. Plates were incubated overnight at 4 °C. Before proceeding with the assay, wells were blocked with 2% bovine serum albumin (BSA; Sigma-Aldrich) for 1 h at room temperature. CPS was detected using serotype-specific rabbit antisera (Statens Serum Institute, Copenhagen, Denmark). After three washes with PBS containing 0.5% Tween-20 (PBST), 100 μL of primary antibody diluted 1:1,000 in BSA-PBS was added to each well and incubated for 2 h at room temperature. Plates were then incubated with alkaline phosphatase-conjugated goat anti-rabbit IgG (Sigma-Aldrich) at a 1:4,000 dilution for 1.5 hours, followed by additional PBST washes. The reaction was developed using p-nitrophenyl phosphate substrate (Sigma-Aldrich), and absorbance was measured at 405 nm after a 30–60 min incubation in the dark at room temperature.

**Mouse model of pneumococcal disease.** Female and male 4-5week old Balb/c mice (Charles River Laboratories) were infected intranasally with $10^7$ cfu/ml Spn resuspended in 50 μl PBS. The mock infection control group received 50 μl of PBS. Some mice were infected intraperitoneally with $10^7$ CFU/ ml Spn resuspended in 100 μl PBS, while the control group received 100 μl PBS. Prior to the infection, the animals were anesthetized via inhalation of 5% isoflurane. Survival was monitored for up to four days. To avoid cross bacterial infection, mice with the same strain where housed within the same box. Animals were weighted daily whereas their behavior and appearance were monitored two times per day. Mice were euthanized when they lost ≥15% of their body weight, compared to their body weight before infection, or when mice were non-responsive to manual stimulation, and/or if they show signs of illness such as ruffled fur, intermittent hunching, and exhibiting respiratory distress. Blood was collected under anesthesia, and the nasopharyngeal tissue, and lungs, were excised post-euthanasia.

An aliquot of blood was immediately added with THY broth added with 10% glycerol and used for dilution and plating onto BAP containing gentamicin. Another aliquot of blood was left untreated and kept at 4°C for 20 min after which serum was separated by centrifugation at 1000 x g for 20 min. Serum was frozen at -80°C until used. Nasopharyngeal tissue, or the lungs, was split in two aliquots, one of them was fixed with 4% paraformaldehyde (PFA) and immediately stored at -80°C for histological analysis and confocal microscopy. The second aliquot was placed in THY added with 10% glycerol (THY-G) and a cocktail of 1X protease inhibitors and immediately homogenized after which the specimens were stored at -80°C. This aliquot was utilized for bacterial counts and for Western blot.

**Depletion of complement.** For survival experiments, mice were treated with 20 µg of cobra venom factor (CoVF) from Naja naja kaouthia (Sigma Cat #8406, 0.8 mg solid with 0.25 mg protein) diluted in 100 µl of saline solution, or with a saline control. The CoVF was administered via three intraperitoneal (i.p.) injections at 12-hour intervals. The Spn strains lacking $H_2O_2$ were intranasally administered at $10^7$ CFU/ml in 50 µl PBS, while the control group received only 50 µl PBS. All animals were sacrificed 4 days post-Spn infection.

**Histopathology**. The PFA-fixed lungs were paraffin-embedded, sectioned (~5 µm) and stained with hematoxylin and eosin (H&E) at the UMMC histology core facility. Micrographs of the tissues were captured with an Olympus BX63 inverted fluorescence microscope. The evaluation of cytotoxicity in various cellular compartments of the lung was conducted blindly by a certified Pathologist. Inflammatory cellular infiltration, bronchiolar hypertrophy, and the extent of bronchiolar exudate and cellular infiltration were assessed using a five-point severity scoring method.[11]

**Quantification of immunofluorescence images.** Confocal z-stacks micrographs were analyzed using the NIS Elements Basic Research software, version 4.30.01 build 1021. For quantifying encapsulated and non-encapsulated bacteria in lung of infected mice, a similar method was used as detailed above for the quantification of encapsulated bacteria in inocula. Region of interest (ROI) was selected to quantify bacteria in the different compartments of the lung. For 3D visualization and quantification purposes, images were processed using the Imaris software 64x Version 10.1.0 (Oxford Instruments). To quantify capsule production by pneumococci, capsule or DNA was identified as a single spot of fluorescence (dot). The software identified these dots and an algorithm calculates their number and percentage of pneumococci (DNA positive bacteria) expressing capsule. Additionally, Imaris was used to identify interactions between immune cells and pneumococci. The area of the cell mask was determined by Imaris, and the square of the area was calculated automatically.

**Whole-genome sequencing.** Genomic DNA was purified using the Qiagen QIAamp DNA Mini Kit as instructed by the manufacturer. The DNA was sequenced at the SeqCenter (Before known as MiGS Microbial Genome Sequencing Center) using the NextSeq 2000 platform. The paired-end read data were assembled and annotated using tools available on RAPT NCBI using g reference strain TIGR4 (AE005672). The sequences were deposited on NCBI GenBank (PRJNA1018957).

**Preparation of a TIGR4Δ*spxB*Δ*lctO* expressing the green fluorescence protein (GFP).** The gene encoding GFP, including a downstream chloramphenicol resistant cassette (*cat*), was amplified from a strain MK147[12], and the fragment was transformed into competent cells of TIGR4Δ*spxB*Δ*lctO*. Chloramphenicol resistant colonies were selected in BAP containing 3.5 µg/ml of the antibiotic. The insertion of the *gfp-cat* fragment was confirmed by PCR and the expression of the GFP was verified by fluorescence microscopy.

**RNA extraction and qRT-PCR analysis.** Cells infected with Spn in 6-well plate collected after 4, 8, 12, and 16 h. Then cells were harvested, and the total RNA was extracted using the RNeasy Plus Mini Kit (QIAGEN) following the manufacturer's instructions. DNA was removed using the Turbo DNA-free Kit (Ambion by Life Technologies). The RNA concentration was obtained using a NanoDrop spectrophotometer (Thermo Fisher Scientific), and 1000 ng of RNA was cDNA transcribed using the iScript cDNA Synthesis Kit (Bio-Rad). Gene expression analysis was carried out using PerfeCTa qPCR ToughMix (QuantaBio) with ready to use mixture of primers with FAM probe (TaqMan® Gene Expression Assays, Thermo Fisher Scientific) and a CFX96 Touch real-time PCR system (Bio-Rad). The primers used for the qRT-PCR analysis were GAPDH, Cat, Nrf1, SOD1, and HO-1. The following conditions were utilized: 1 cycle at 95°C for 3 min and 40 cycles of 95°C for 15 s, 60°C for 30 s and 72°C for 30 s. Melting curves were generated to confirm the absence of primer dimers. The relative quantitation of mRNA expression was normalized to the constitutive expression of the housekeeping GAPDH gene and calculated by the comparative cycle threshold (2−ΔΔCT) method.

Spn was inoculated in THY broth with 1% sheep red blood cells (sRBC), THY containing Hb (1 or 10 µM), or THY alone. Pneumococci were also inoculated into cultures of A549, or Calu-3 cells, added with cell culture medium supplemented with 1% sRBC. These cultures were incubated for 2 h at 37 °C in a 5% $CO_2$ atmosphere. Bacteria were then collected, RNA extracted, and qRT-PCR analysis were performed as described by our group earlier[13] using primer targeting the *spxb* or the *cps4A* gene. The relative quantitation of mRNA expression was normalized to the constitutive expression of the housekeeping 16S rRNA gene and calculated using the comparative CT ($2^{-\Delta\Delta CT}$) method.

**RNA-seq analyses.** The datasets deposited in NCBI (accession IDs: PRJNA626052[1], PRJNA642413[2], and PRJNA637100[3]), were utilized to investigate the expression of genes in the D39 background in response to different culture conditions and infection conditions. The *in vitro* dataset, labeled as bioproject PRJNA626052, compares RNA-seq data with or without Hb conditions, while PRJNA642413 consists of expression profiles of bacterial biofilm and planktonic cells in the presence of Hb. For *in vivo* dual RNA-seq data, we utilized bioproject PRJNA637100, which provides dual RNA-seq profiles obtained from infected mice from 2-7 days at disease-relevant anatomical sites, as well as from blood, identifying organ-specific transcriptomes.

The original dataset of PRJNA626052[1] was analyzed by the Molecular and Genomics Core Facility at the University of Mississippi Medical Center. Differential expression of bacterial genes was assessed between different culture conditions, containing or not (controls) 20 µM Hb. Quality control steps were performed on raw datasets. Reads were aligned to the NCBI Refseq genome D39 using the basespace application RNA-Seq Alignment (Version: 2.0.1), which conducted splice-aware genome alignment with STAR (version 2.6.1a). Differential expression (DE) analysis of genes (DGE) was performed using DESeq2 (version 1.20.0).

The gene encoding enzymes of the pyruvate metabolic network and capsule genes were assessed to determine which genes were overexpressed across the context-relevant datasets. The selected genes were sorted based on their average fold-change and TPM (transcripts per million) across these datasets and were visualized using a heatmap generated by GraphPrism v10.1.

**Primary metabolism analysis of pneumococcal cultures containing oxidized hemoglobin**. THY or THY containing 10 µM hemoglobin (Hb-$Fe^{3+}$), was infected with either TIGR4 or TIGR4Δ*sxB*Δ*lctO,* in a 6-well microplate, and the cultures were incubated for 2 or 4 h at 37°C in a 5% $CO_2$ atmosphere. The culture supernatant was removed, filter sterilized with a 0.22 µm syringe filter and immediately frozen at -80°C. Aliquots of these frozen supernatants were analyzed by spectroscopy to investigate the oxidation of hemoglobin. Primary metabolism by gas chromatography-time of flight mass spectrometry (GC-TOF MS) was performed at the West Coast Metabolomics Center, UC Davis. Briefly, samples extracted using 1ml of 3:3:2 ACN:IPA:$H_2O$ (v/v/v). Half of the sample was dried to completeness and then derivatized using 10 µl of 40 mg/ml of methoxyamine in pyridine. They were shaken at 30°C for 1.5 h. Then 91 µl of MSTFA + FAMEs to each sample and they were shaken at 37°C for 0.5 h to finish derivatization. Samples were then vialed, capped, and injected onto the instrument. We use a 7890A GC coupled with a LECO TOF. 0.5 µl of derivatized sample is injected using a splitless method onto a RESTEK RTX-5SIL MS column with an Intergra-Guard at 275°C with a helium flow of 1 ml/min. The GC oven is set to hold at 50°C for 1 min then ramp to 20°C/min to 330°C and then hold for 5 min. The transferline was set to 280°C while the EI ion source was set to 250°C. The Mass spec parameters collect data from 85m/z to 500m/z at an acquisition rate of 17 spectra/sec. Raw data were processed by LECO ChromaTOF version 4.5 for baseline subtraction, deconvolution and peak detection, while BinBase was used for annotation and reporting.[14] GC-TOF MS data were deposited in the Metabolomics Workbench repository (Study ID: STXXXXX), accessible at https://www.metabolomicsworkbench.org.

**Quantification of metals in lung tissue.** Elemental images of lung sections were collected via laser ablation inductively coupled plasma time of flight mass spectroscopy (LA-ICP-TOF-MS) at the Biomedical National Elemental Imaging Resource (BNEIR) at Dartmouth College (NIGMS R24GM141194) in April 2023. Instrumentation consisted of an imageBIO266 laser ablation unit (Elemental Scientific Lasers,) and a Vitesse ICP-TOF-MS detection system (Nu Instruments,).

Laser ablation used the imaging cup, a 10 µm spot size, 150 Hz repetition rate, with He gas flow through both imaging cup and sample chamber at 225 L/min, a scan speed of 1500 µm/sec and a fluence of 3 $J/cm^2$. The LA unit was connected to the Vitesse via a Dual Concentric Injection (DCI) connection unit, with a nebulizer argon flow of 1150 L/min, and the Vitesse was operated in $H_2$ gas mode. The mass range scanned was 22 to 260, omitting masses 27.5 to 40.5 to prevent over-saturation of the Vitesse detector. The LA-ICP-TOF-MS was tuned daily using the NIST 612 glass standard to optimize performance.

Quantification of detector counts into parts per million used analysis of a custom-made micro-spotted gel standard calibration series[15] consisting of 5 concentration levels, collected before and after analysis of each sample. Data reduction was conducted using the Iolite application package[16], which conducts background subtraction and quantification using the Howell method [17] as part of the 3D Trace Elements data reduction scheme.

**Statistical analysis.** The statistical analysis of primary metabolism involved normalizing to the median intensity of all identified compounds, applying log transformation, and using Pareto scaling. Principal Component Analysis (PCA) was utilized for multivariate statistics and visualization, particularly for detecting outliers. Results from Kruskal-Wallis tests were followed by Dunn's multiple comparison adjustment. Results from Mann-Whitney U tests were corrected using the Benjamini-Hochberg procedure to control the false discovery rate. Spearman rank correlation analyses and fold change calculations were performed using R. We performed one-way analysis of variance (ANOVA) followed by Dunnett's multiple-comparison test when more than two groups were compared or Student's t test to compare two groups, as indicated. All statistical analysis was performed using the software GraphPad Prism (version 8.3.1).

**Table S1. Metabolites differentially regulated in the supernatant of THY-$Hb^{3+}$ cultures infected with TIGR4 compared to plain THY.**

| Molecule | Class | Adjusted *p*-value | Fold-change |
|---|---|---|---|
| **THY-$Hb^{3+}$/TIGR4 compared with THY/TIGR4** | | | |
| Aspartic acid | Organic acid | 0.006 | 0.955886 |
| Creatinine | Organic acid | 0.003 | 1.616839 |
| Guanine | Organoheterocyclic compounds | 0.026 | 0.551192 |
| Guanosine | Nucleosides, nucleotides, and analogues | 0.026 | 1.301212 |
| Isoleucine | Organic acids and derivatives | 0.013 | 1.051325 |
| Leucine | Organic acids and derivatives | 0.013 | 1.079021 |
| Methionine | Organic acids and derivatives | 0.004 | 0.591517 |
| Methionine sulfoxide | Organic acids and derivatives | 0.004 | 2.001035 |
| Nicotinamide | Organoheterocyclic compounds | 0.031 | 0.899909 |
| Oleamide NIST | Lipids and lipid-like molecules | 0.006 | 1.119466 |
| Oxoproline | Organic acids and derivatives | 0.006 | 0.945675 |
| Putrescine | Organic nitrogen compounds | 0.004 | 1.683839 |
| Pyruvic acid | Organic acids and derivatives | 0.004 | 0.053013 |
| Shikimic acid | Organic oxygen compounds | 0.003 | 0.722413 |
| Tyrosine | Organic acid | 0.013 | 1.077018 |
| Tyrosol | Benzenoids | 0.004 | 1.614024 |
| **THY-$Hb^{3+}$/TIGR4Δ*spxB*Δ*lctO* compared with THY/TIGR4Δ*spxB*Δ*lctO*** | | | |
| Creatinine | Organic acid | 0.003 | 2.320286 |
| Shikimic acid | Organic oxygen compounds | 0.003 | 0.623841 |
| **THY-$Hb^{3+}$/TIGR4 compared with THY-$Hb^{3+}$/TIGR4Δ*spxB*Δ*lctO*** | | | |
| Creatinine | Organic acids and derivatives | 0.01 | 1.457524 |
| Nicotinamide | Organoheterocyclic compounds | 0.003 | 1.553198 |
| Nicotinic acid | Organoheterocyclic compounds | 0.003 | 0.753161 |
| Tryptophan | Organoheterocyclic compounds | 0.026 | 0.923314 |
| Tyrosine | Organic acids and derivatives | 0.013 | 0.88953 |
| Tyrosol | Benzenoids | 0.004 | 0.589645 |

| | | | |
|---|---|---|---|
| **THY/TIGR4 compared with THY/TIGR4Δ*spxB*Δ*lctO*** | | | |
| Guanosine | Nucleosides, nucleotides, and analogues | 0.026 | 1.204739 |
| Methionine | Organic acids and derivatives | 0.004 | 0.57766 |
| Methionine sulfoxide | Organic acids and derivatives | 0.004 | 2.064243 |
| N-acetylornithine | Organic acids and derivatives | 0.013 | 0.840103 |
| Nicotinamide | Organoheterocyclic compounds | 0.003 | 1.692531 |
| Nicotinic acid | Organoheterocyclic compounds | 0.003 | 0.762097 |
| Oleamide NIST | Lipids and lipid-like molecules | 0.017 | 1.082949 |
| Orotic acid | Organoheterocyclic compounds | 0.013 | 0.667873 |
| Putrescine | Organic nitrogen compounds | 0.004 | 1.622776 |
| Pyruvic acid | Organic acids and derivatives | 0.004 | 0.178594 |
| Shikimic acid | Organic oxygen compounds | 0.031 | 1.068225 |

**Table S2. Pneumococcal strains utilized in this study.**

| Strain | Characteristics | Reference |
|---|---|---|
| TIGR4 | Clinical isolate, serotype 4 | Tettelin et al., 2001 |
| P2422 | TIGR4Δ*cps*; Kan$^R$ | Zafar et al., 2016 |
| SPJV29 | TIGR4Δ*spxB*; Ery$^R$ | McDevitt et al., 2020 |
| SPJV41 | TIGR4Δ*spxB*Δ*lctO*; Ery$^R$, Spc$^R$ | McDevitt et al., 2020 |
| SPJV61 | TIGR4Δ*spxB*Δ*lctO*Ω*gfp*; Ery$^R$, Spc$^R$, Cam$^R$ | This study |
| SPJV44 | TIGR4Δ*spxB*Δ*lctO*Ω*spxB-lctO*; Ery$^R$, Spc$^R$, Cam$^R$ | Alibayov et al., 2022 |
| EF3030 | Clinical isolate, serotype 19F | Petersen et al., 2019 |
| SPJV49 | EF3030Δ*spxB*Δ*lctO*; Ery$^R$, Spc$^R$ | McDevitt et al., 2020 |
| D39 | clinical isolate, serotype 2 | Avery et al., 1944 |
| SPJV45 | D39Δ*spxB*Δ*lctO*; Ery$^R$, Spc$^R$ | McDevitt et al., 2020 |
| R6 | D39 derivative serotype 2 (smooth or encapsulated) | Avery et al., 1944 |
| SPJV55 | R6Δ*spxB*Δ*lctO*; Ery$^R$, Spc$^R$ | McDevitt et al., 2020 |

**Supplementary Figures**

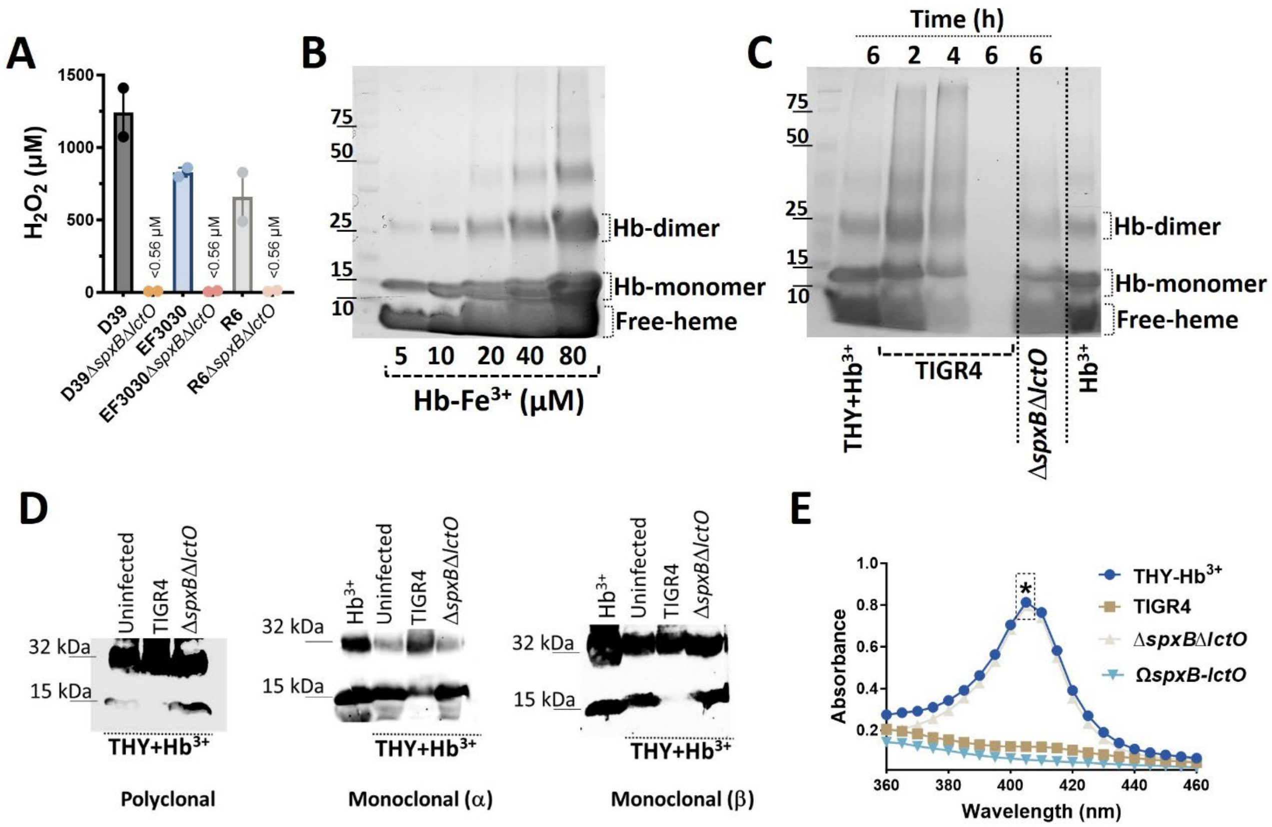

**Supplementary Fig. 1**. **Oxidation of hemoglobin by Spn-$H_2O_2$ leads to heme degradation and hemoglobin polymerization.** (A) Bacterial strains D39, D39Δ*spxB*Δ*lctO*, EF3030, EF3030Δ*spxB*Δ*lctO*, R6, or R6Δ*spxB*Δ*lctO* were grown in THY media at 37°C for 6 hours. Supernatants were collected, and $H_2O_2$ levels were quantified using Amplex Red (n=2 independent experiments, LOD < 0.56 µM). (B) Native PAGE analysis of THY media with increasing concentrations of Hb-$Fe^{3+}$. Heme staining visualized the presence of heme. Molecular weights (KDa) are indicated on the left. (Hb=hemoglobin). THY media with 10 µM $Hb^{3+}$ (THY-$Hb^{3+}$) was incubated with or without pneumococcal infection for various times (C). Supernatants were analyzed by native PAGE with $Hb^{3+}$ (10 µM) as a control and heme was stained. Representative experiments from ≥4 independent replicates. (D) THY-$Hb^{3+}$ (20 µM) was incubated with TIGR4, TIGR4Δ*spxB*Δ*lctO*, or remained uninfected for 6 hours. Supernatants were analyzed by Western blot using polyclonal anti-hemoglobin, anti-α hemoglobin, or anti-β hemoglobin antibodies . $Hb^{3+}$ standard was loaded as a control. (E) Spectroscopy analysis of supernatants from cultures in (D). Absorbance spectra (360-460 nm) were collected. Student's t-test compared absorbance at 405 nm between TIGR4Δ*spxB*Δ*lctO* and TIGR4 or TIGR4Ω*spxB-lctO* cultures (*$p<0.05$).

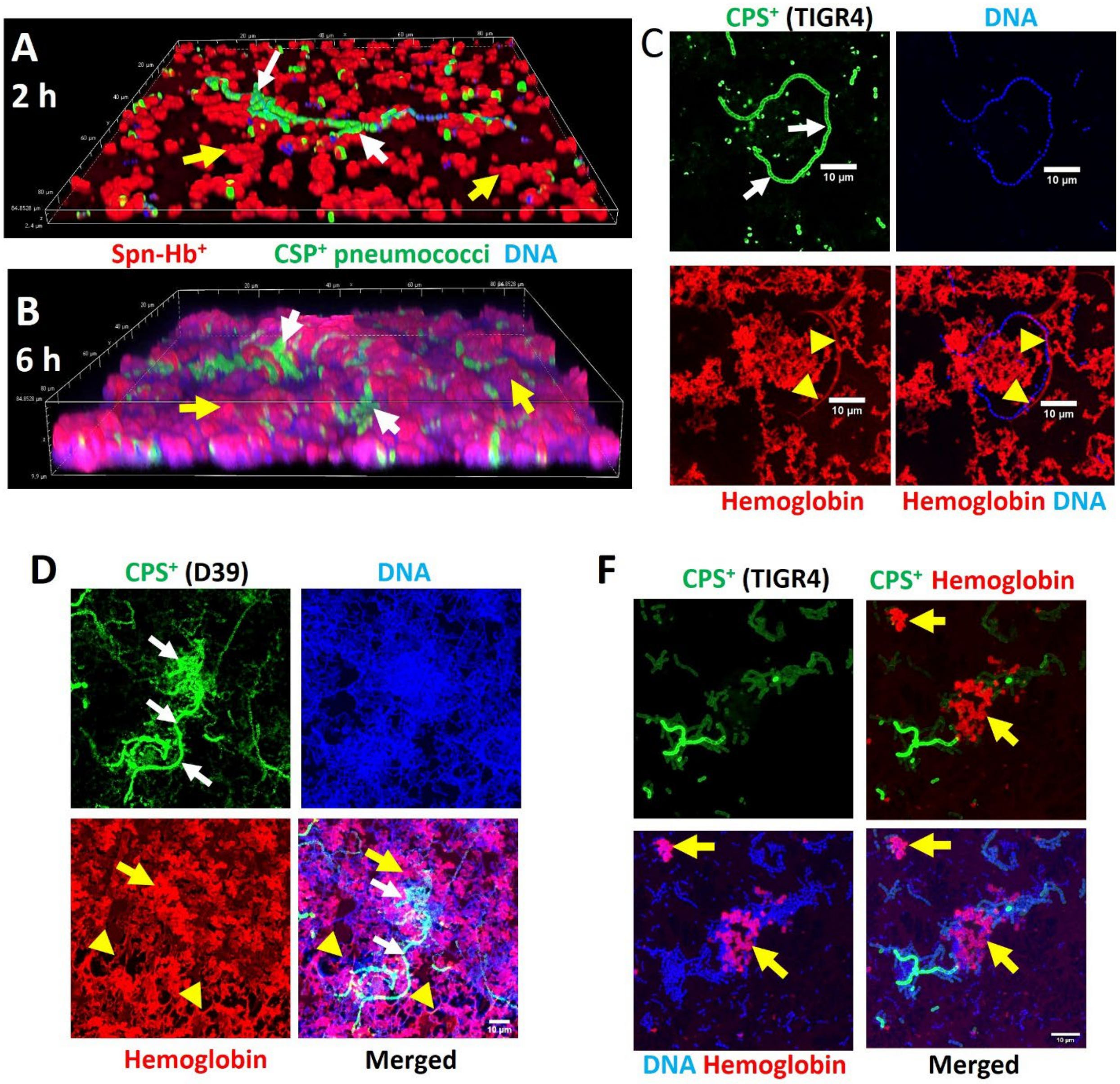

**Supplementary Fig. 2**. **Hemoglobin oxidation selects for encapsulated pneumococci**. TIGR4 (all panels except D) or D39 (D) was cultured in THY-Hb$^{3+}$ (10 µM) for 2 (A) or 6 hours (B-F) at 37°C with 5% $CO_2$. Biofilm (A-D) and planktonic bacteria (F) were stained with anti-Spn capsule and anti-hemoglobin antibodies, and DNA was counterstained with DAPI. Confocal microscopy images represent at least three independent experiments. Panels A and B show 3D reconstructions from 0.5 µm z-stacks. White arrows indicate CPS$^+$ pneumococci, yellow arrows indicate Spn-Hb$^+$ bacteria, and arrowheads indicate hemoglobin aggregates.

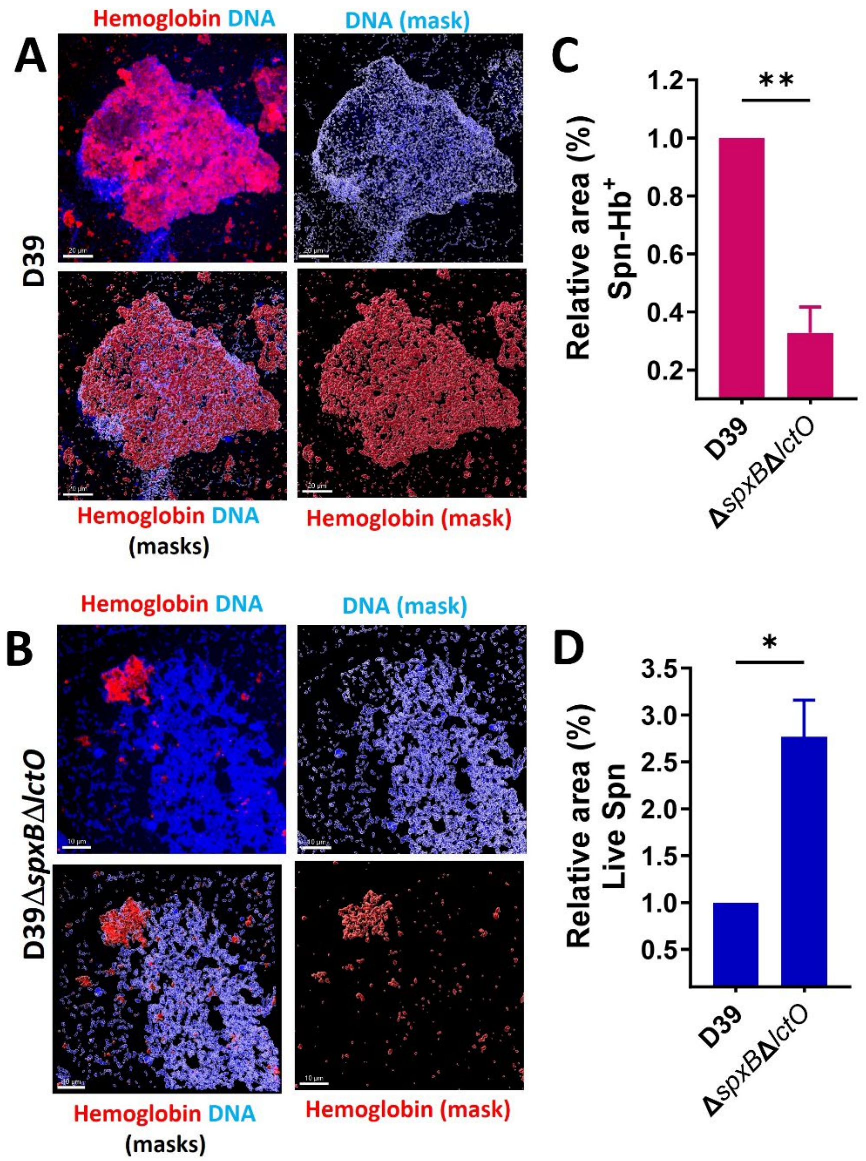

**Supplementary Fig. 3**. **Hemoglobin oxidation intoxicates planktonic pneumococci.** D39 (A) or D39Δ*spxB*Δ*lctO* (B) pneumococci were incubated with 10 µM THY-Hb$^{3+}$ for 6 hours at 37°C. Planktonic bacteria were stained with anti-hemoglobin antibody and DAPI for DNA visualization. Confocal microscopy was used to acquire images. Shown in (A) and (B) are projections of z-stacks (top left panels). Imaris software was used to create masks for each channel, allowing quantification of Spn-Hb$^{+}$ pneumococci (C) or live pneumococci (D) in D39Δ*spxB*Δ*lctO* relative to D39. Data represent mean ± SEM from two independent experiments analyzing at least ~20,000 pneumococci per strain. Statistical significance was determined using a Mann-Whitney test (*p < 0.05, **p < 0.001).

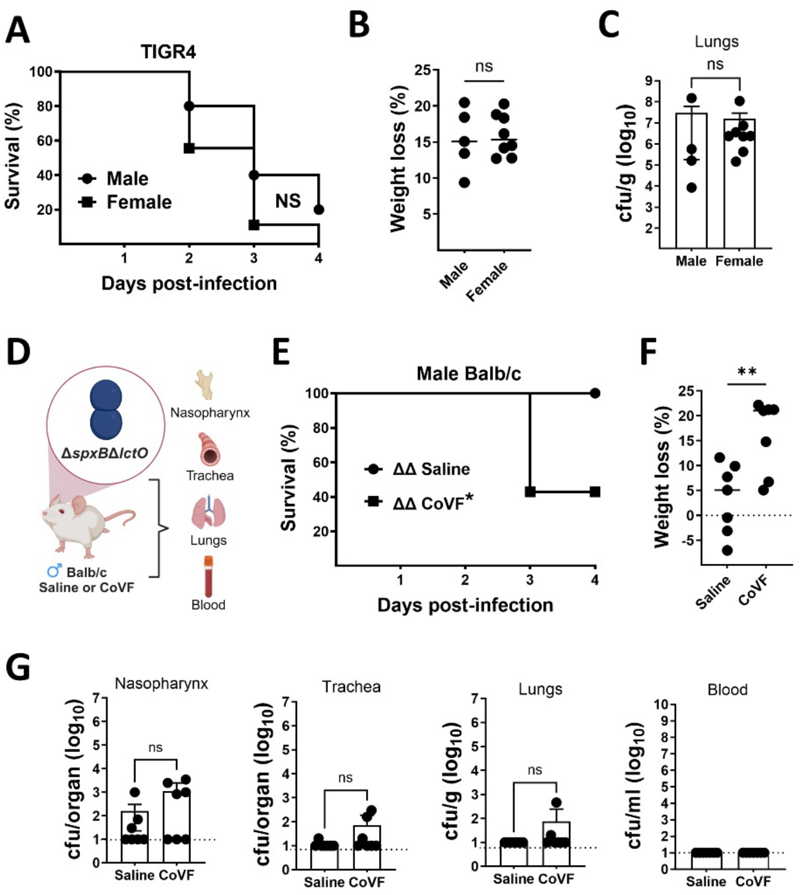

**Supplementary Fig. 4**. **Increase survival of Balb/c mice infected with TIGR4 lacking production of $H_2O_2$.** (A-C) Female (n=8) or male (N=5) Balb/c mice received ~$10^7$ CFU TIGR4 intranasally. (A) Mortality was monitored for four days and (B) weight loss at euthanasia relative to pre-infection weight is shown. (C) Bacterial burden was determined in the lungs. (D) Schematic of treatment, saline or cobra venom factor (CoVF), intranasal inoculation with ~$10^7$ CFU TIGR4Δ*spxB*Δ*lctO* and harvested organs at euthanasia. (E) Mortality was monitored for four days and (F) weight loss at euthanasia relative to pre-infection weight is shown. Bacterial burden was determined in (G) nasopharynx, trachea, lung, and blood. Log-rank Mantel-Cox test for survival (A and E): NS=not significant, **p<0.005 vs. saline/TIGR4Δ*spxB*Δ*lctO*. Unpaired student's *t* test was performed in panels (B, C, F-G) NS=not significant, **p<0.005.

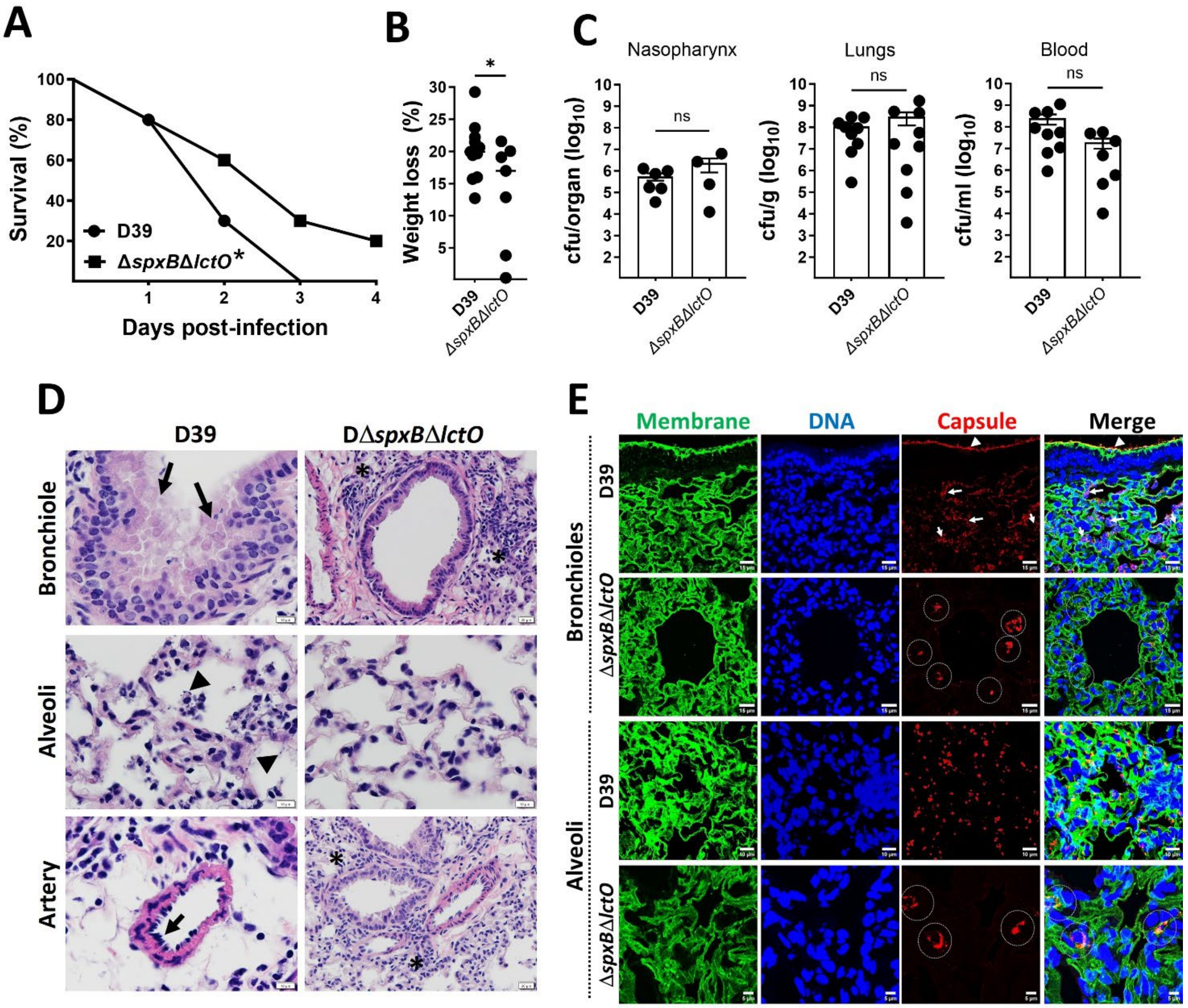

**Supplementary Fig. 5**. **Increase survival of mice infected with Spn D39 strain lacking production of $H_2O_2$.** (A) Balb/c mice (n=12) were intranasally inoculated with ~$10^{7}$ CFU of D39, while a control group (n=8) received D39Δ*spxB*Δ*lctO*. Survival was monitored for four days and analyzed by log-rank Mantel-Cox test (*p<0.05). (B) Mice were weighed daily, and percent weight loss at euthanasia relative to pre-infection weight is shown. (C) Bacterial burden was quantified in nasopharynx, lung, and blood. Unpaired t-test was used for statistical analysis (*p<0.05, **p<0.005, ***p<0.0005). (D) Lung sections were stained with hematoxylin and eosin. Micrographs depict bronchioles, alveoli, and arteries. Arrows indicate bronchiolar or endothelial cell toxicity, arrowheads denote pneumococci, and asterisks mark areas of inflammatory infiltration. (E) Lung sections from Spn-infected mice were immunostained for membrane, capsule, and DNA. Encapsulated D39 bacteria colonized bronchioles (arrowheads) and invaded alveoli (arrows). D39Δ*spxB*Δ*lctO* bacteria formed aggregates in both bronchioles and alveoli (circled). Confocal microscopy z-stack projections of bronchioles and alveoli are shown.

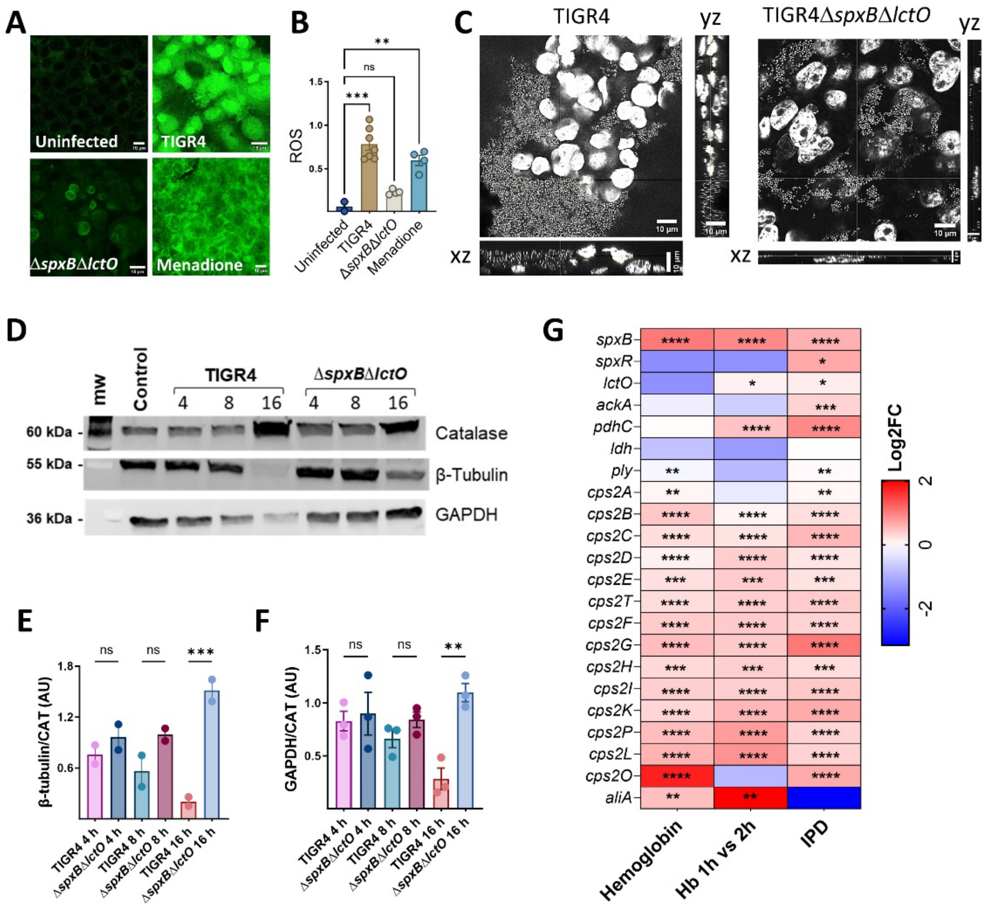

**Supplementary Fig. 6**. **Intracellular Spn causes an increase in levels of reactive oxygen species (ROS) in A549 cells and triggers degradation of cytosolic proteins.** (A-C) Human alveolar A549 cells were exposed to TIGR4 or TIGR4Δ*spxB*Δ*lctO*, supplemented with 10 µM menadione, or left untreated for 8 h at 37°C with 5% $CO_2$. Subsequently, cells were labeled with (A) CellROX (5 µM) for 30 min, fixed, and counterstained with DAPI. (B) Confocal microscopy and z-stack projection were employed for ROS quantification. (C) Representative confocal XY, XZ, and YZ planes from DAPI channel are shown. (D) Catalase, β-tubulin, and GAPDH protein levels were assessed by Western blot in mock- or Spn-infected A549 cells at indicated timepoints in hours. (E-F) β-tubulin or GAPDH protein band intensities were normalized to catalase and expressed as arbitrary units. Data represent mean ± SEM (n ≥ 2). (G) Heatmap visualization compares gene expression levels. Genes analyzed include those encoding pyruvate node enzymes, pneumolysin (*ply*) and capsule locus genes. Samples are from D39 cultures grown on THY-Hb$^{3+}$ (Hemoglobin) vs. THY, THY-Hb$^{3+}$ incubated for 2 h vs. 1 h and from *In vivo* TIGR4-infection model of invasive pneumococcal disease (IPD) compared with gene expression in the nasopharynx.

Statistical significance was determined by (B, E-F) one-way ANOVA with Dunnett's post hoc test or (G) the non-parametric Wilcox test; *p<0.05, **p<0.01, ***p<0.0005, ****p<0.0001, ns= not significant.

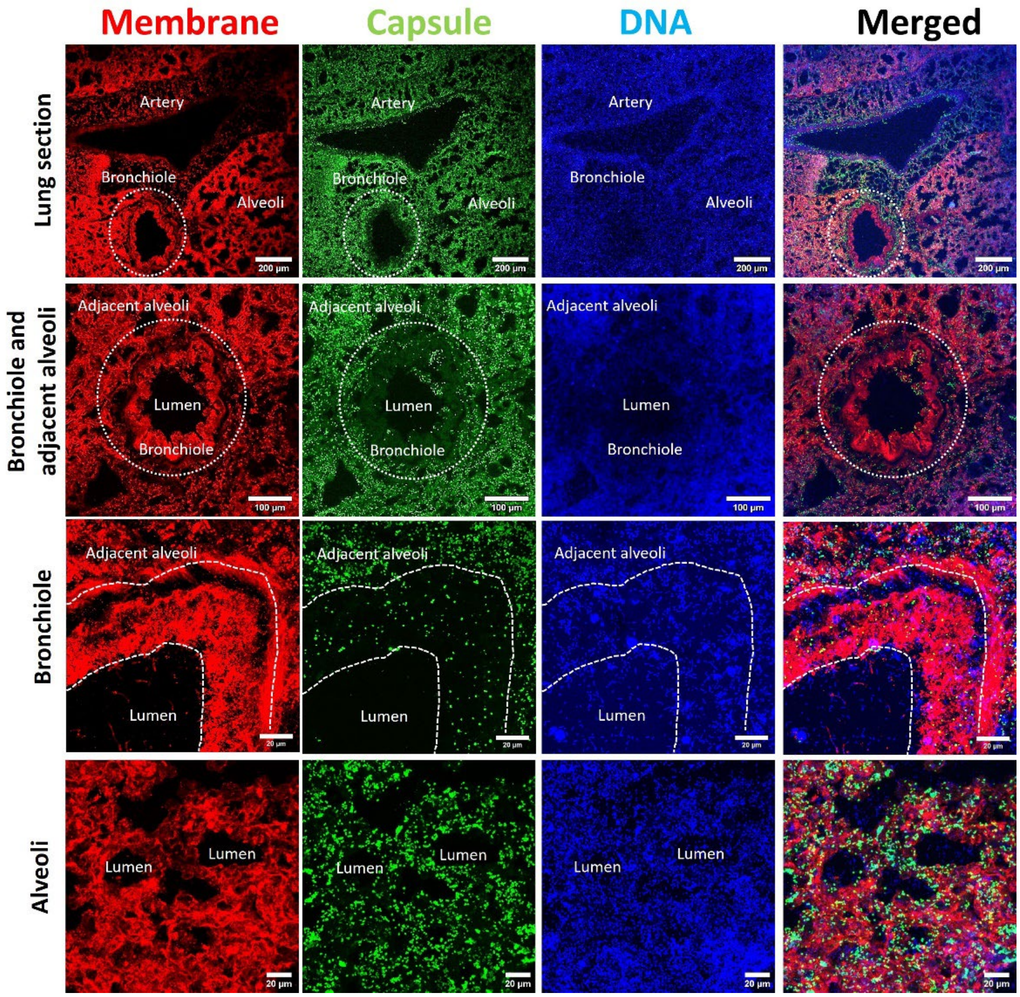

**Supplementary Fig. 7. Spn encapsulates in the alveolar epithelium in a mouse model of pneumococcal pneumonia.** Balb/c mice received a single intranasal dose of TIGR4 (~$10^7$ CFU) and were euthanized upon onset of terminal disease. Lungs were aseptically harvested, processed, and stained for cell membranes, capsule, and DNA. Confocal microscopy was employed to image bronchioles, alveoli, or pulmonary arteries, with data presented as z-stack projections. A region of interest was demarcated (dashed line) encompassing the non-encapsulated bacterial population within the pseudostratified bronchiolar epithelium, excluding adjacent alveoli characterized by pneumococcal re-encapsulation.

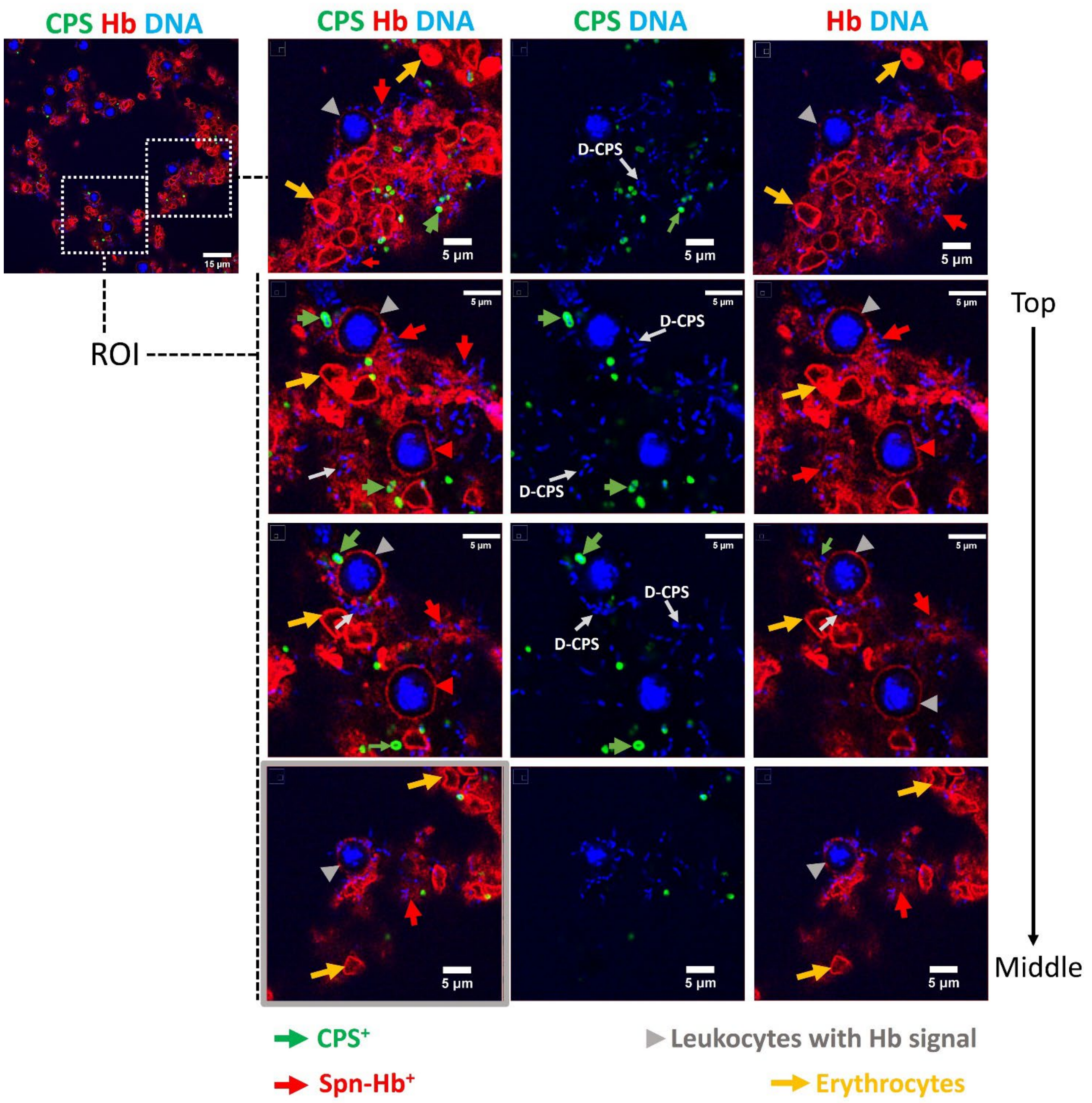

**Supplementary Fig. 8**. **Spn causes the polymerization of hemoglobin in the alveoli of mice with pneumococcal pneumonia.** Balb/c mice infected intranasally with TIGR4 were euthanized upon moribundity. Lungs were aseptically harvested, fixed, embedded in paraffin, and sectioned (5 µm). Sections were immunostained for capsule ($CPS^+$), hemoglobin (Hb), and DAPI was used to stain DNA. Alveolar epithelium was imaged by confocal microscopy, generating 0.3 µm XY optical sections. The left panel displays a projection, while the right panels show optical sections of the indicated zoomed region. Green arrows: encapsulated ($CPS^+$) pneumococci; red arrowheads: leukocytes with Hb signal; grey arrows: Spn-$Hb^+$ bacteria; yellow arrows: erythrocytes; white arrows: D-CPS.

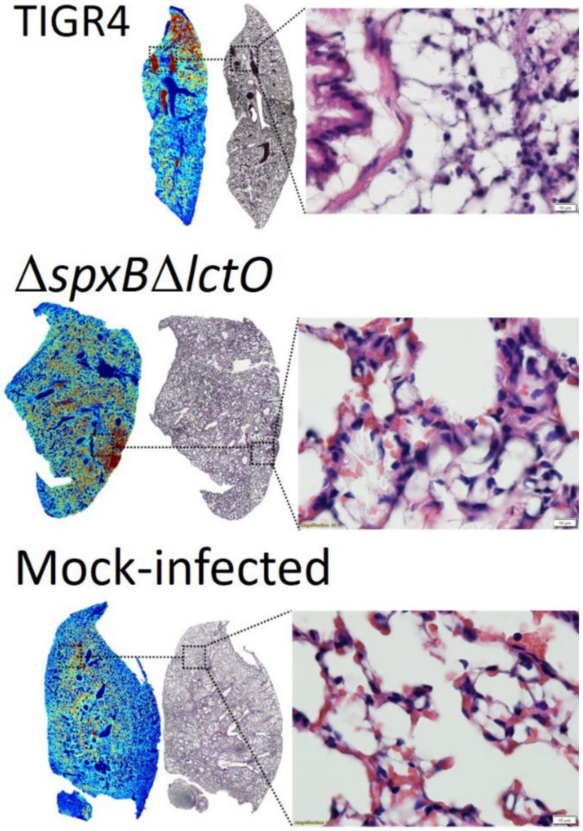

**Figure S9. Analysis of the spatial localization of iron in the lung of mocked-infect, and mice infected with pneumococci.** Spatial distribution of $^{56}Fe^{+}$ was assessed in mouse lung sections by LA-ICP-MS following infection with (A) TIGR4, (B) TIGR4Δ*spxB*Δ*lctO*, or (C) mock infection. Hematoxylin and eosin staining of corresponding lung sections (middle panels) and higher magnification histology micrographs of selected lung regions are included.

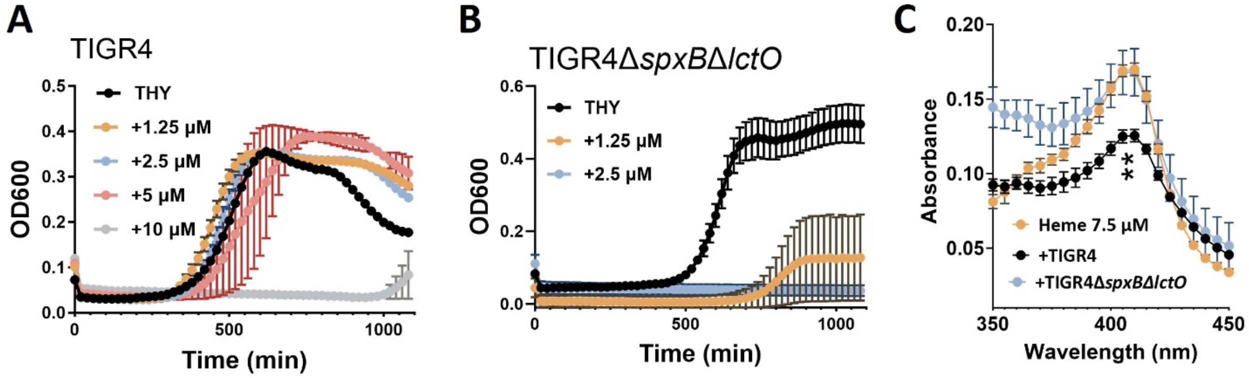

**Supplementary Fig. 10**. **Heme is toxic for pneumococci lacking production of hydrogen peroxide.** (A-B) Growth curves of

TIGR4 and TIGR4Δ*spxB*Δ*lctO* strains cultured in THY medium with or without indicated heme concentrations. Cultures were incubated at 37°C with 5% $CO_2$, and optical density at 600 nm was measured every 20 minutes. (C) Supernatants from THY cultures supplemented with 7.5 µM heme and inoculated with TIGR4, TIGR4Δ*spxB*Δ*lctO*, or left uninfected for four hours were analyzed by spectrophotometry (350-450 nm). Mann-Whitney U test was used to compare Soret peak absorbance (405 nm) between uninfected and pneumococci-infected media (**$p<0.001$).
